# Functional and causal neural mechanisms of human voice perception in noisy situations

**DOI:** 10.1101/2024.10.21.619396

**Authors:** Leonardo Ceravolo, Elisa Scariati Jaussi, Sascha Frühholz, Dimitri Van De Ville, Didier Grandjean

**Affiliations:** Neuroscience of Emotion and Affective Dynamics Lab, Department of Psychology, University of Geneva, Geneva, Switzerland; Swiss Center for Affective Sciences, University of Geneva, Geneva, Switzerland; Geneva University Hospital, Geneva, Switzerland; Department of Psychology, University of Zurich, Zurich, Switzerland; Department of Psychology, University of Oslo, Oslo, Norway; Department of Radiology and Medical Informatics, University of Geneva, Geneva, Switzerland; Institute of Bioengineering, Ecole Polytechnique Fédérale de Lausanne, Lausanne, Switzerland

**Author notes:** **Corresponding author:** Leonardo Ceravolo, University of Geneva & Swiss Center for Affective Sciences, Unimail, office 5133, Boulevard Pont-d’Arve 50, CH-1205 Geneva, Switzerland.

## Abstract

Human communication entails an efficient way of simultaneously processing voice and reducing the impact of environmental noise. By manipulating background noise, we aimed at clarifying the neural mechanisms allowing voice comprehension in noisy situations. Our results point to spatial and temporal coexistence of lateral and medial temporal cortex networks when voice is easily detected in highly noisy conditions, revealing the necessary neural underpinnings of human communication in realistic situations.

---

Among the most recent evolutionary changes that made *Homo sapiens* today’s anatomically modern human^1^, language and vocal communication were of the highest importance and have therefore been extensively studied by the neuroscientific community^2^. Voice processing—independently of semantics—was however in the limelight, especially in the context of cognitive neuroscience, until voice-sensitive areas were identified along the bilateral human temporal lobe^3^. These voice-sensitive areas include large portions of the middle and superior temporal lobe or superior temporal cortex (STC) and underlie vocal processing in anterior, mid and posterior subparts of the STC^4^. Notably, parts of the posterior STC have been shown to process information about voices, faces and face-voice integration in humans^5^, raising the question of whether voice-sensitive brain regions are species-specific or species-independent. Species-specific, voice-sensitive/vocalization-specific areas have been highlighted in studies on the STC of humans^6^ and macaques^7^, notably in the anterior temporal sulcus for the former as well as in the posterior ‘belt’ and ‘parabelt’ regions of the auditory cortex for the latter. In humans, subparts of these species-specific, voice-sensitive areas—in particular, the mid and posterior parts of the STC—respond strongly to general emotional voice content^8-10^ and more specifically to angry vocal signals^8,9,11-16^, extending previously mentioned findings that focused mainly on non-emotional voice perception. The posterior STC was also highlighted as an important bilateral region involved in recognizing environmental sounds^17^ and voice-in-noise perception in adults^18^, in addition to the bilateral posterior middle temporal gyrus (MTG)^19^, emphasizing distinct roles among temporal lobe subregions usually taken as a whole and referred to as ‘voice-sensitive’.

One can however question whether voice-related processing mainly relies on the bilateral STC—and its subparts—as specialized hubs for processing vocal signals, or whether these areas instead work in a coordinated way with other relevant cortical and/or subcortical brain regions known to be involved in (emotional) vocal processing and/or environmental sound recognition. This aspect is of particular interest since our brain should be able at any time to filter noise—at least partially—to favor vocal communication, especially in the context of social interactions. Relevant areas include the inferior frontal cortex^15,17,19^ and subregions of the medial temporal lobe (MTL; i.e., the bilateral hippocampus, amygdala and parahippocampal gyrus) involved for instance in vocal threat perception^20^. In fact, subregions of the amygdala were shown to underlie the automatic (bilateral superficial complex) and explicit (right laterobasal complex) focus of attention on vocal threat^14,20^. Moreover, the role of the hippocampus in vocal and musical affect processing was emphasized through both direct connections to the primary auditory cortex and indirect connections mediated by the parahippocampal gyrus^21^. The amygdala, parahippocampal gyrus and hippocampus were also shown to underlie the processing of familiar as opposed to non-familiar voices and noises presented both visually and auditorily^22^, adding even more weight to the inclusion of MTL regions in the functional neural framework of voice perception in realistic—often noisy—situations.

While environmental noise is, in the context of the present studies, related to voice perception and may seem almost trivial in comparison to the duties and challenges of everyday life as humans, being exposed to noise may actually play a significant role in several health conditions and cognitive impairments. In fact, recent literature abounded for a role of noise in the induction of stress hormones, oxidative stress and ultimately vascular dysfunctions^23^—as recently reviewed for transportation noise pollution^24^—as well as brain impairment and cognitive decline in mice^25^ when exposed to chronic traffic noise. The same goes for humans^26^, together with potential psychological disorders. Therefore, noise can be categorized as a prevalent and important cause of stress, including above-mentioned risks on health. In the context of our studies—and even though we emphasized the importance and causal relation between an exposition to noise and stress, we specifically challenged the ability of human participants to perceive voice and environmental noise both separately and concurrently. We assumed, based on the literature and on known models of voice processing^27^ that parallel brain networks would be involved, while their specific locations and the involved brain areas still remain unclear. Therefore, an integrated brain-level understanding of voice processing in humans in realistic, noisy situations should question and try to pinpoint the potential existence of such parallelization. This investigation should have as a starting point the different subparts of voice-sensitive areas. In fact, these subparts could act in parallel and/or together with relevant cortical and subcortical brain regions such as the MTL or the lateral/inferior frontal cortex. Identifying and testing such brain network could therefore have the potential to clarify the neural mechanisms allowing noise-free vocal communication. Such questions can be addressed scientifically by the use of static^28^ and dynamic or causal^29,30^ connectivity measures at the brain level. The literature on structural^31^ and functional^4,15,32^ connectivity of voice-sensitive brain areas is however scarce and the lack of evidence for a distributed brain network underlying the full neural spectrum of vocal processing, especially in realistic noisy situations^33^, is needed. The abovementioned research question led us to several assumptions and hypotheses that were operationalized through two independent functional MRI studies and one behavioral study. When voice and noise were presented separately (Study 1, N=98), we hypothesized: (a) enhanced activity and functional and effective connectivity in voice-sensitive areas (STC)—as a replication of existing data3— and MTL^21^, (b) an involvement of MTL and posterior STC when processing environmental sounds and noise^19^. When voice and noise were presented concurrently at various intensities (Study 2, N=18), we predicted: (c) response patterns pertaining to an ease of perceiving a vocal signal shifting towards chance when noise was the highest, (d) with slower reaction times due to an increase of task difficulty (high noise levels). For the same paradigm using fMRI (Study 3, N=20), we hypothesized: (e) a replication of the patterns of behavioral responses found in Study 2, (f) enhanced activity in and functional connectivity between posterior STC and the MTL—specifically the amygdala, parahippocampal gyrus and hippocampus—when processing and assessing voices while background noise was present, especially when noise is at higher acoustic intensities than voices and finally (g) a causal implication of posterior STC, especially the posterior middle temporal gyrus (pMTG) and subregions of the MTL (amygdala, parahippocampal gyrus and hippocampus) for successful vocal communication in noisy situations. The operationalization of our hypotheses should therefore reveal how specific, potentially parallel brain networks may underlie successful and reliable voice processing and comprehension in noisy environments and bring new insights to voice perception and communication in humans. This approach is meant to point out a novel framework targeting the general, coordinated and distributed neural substrates underlying vocal processing rather than focusing solely on temporal voice areas (TVAs) or on auditory brain regions.

## Results

The three studies presented in this manuscript were planned as a grouped effort to approach a better understanding of voice perception in realistic situations—i.e., when environmental noise is omnipresent. The final outcome is illustrated by a general model of voice processing in everyday situations. Study 1 aimed at replicating previous findings^3^ involving ‘voice-sensitive areas’ and also extending our knowledge of how humans process both voice and noise signals, separately. Study 2 was implemented to assess the ability of human participants to process and perceive voice when background noise is present—at varying intensity levels. It was also the starting point to allow an implementation of such paradigm in the MRI scanner (Study 3). Study 3 was therefore acquired to better understand brain networks—both using wholebrain, functional and effective connectivity analyses—allowing a reliable perception of voice signals in noisy situations and the subsequent decision(s) based on such prior processing, which parallels our everyday experience as humans considering social interactions and communication.

### Study 1: voice-sensitive areas and ‘non-vocal’, environmental noise processing

We started with whole-brain analyses in order to replicate the findings of Belin and colleagues^3^ and, of more importance, to have a strong statistical basis to extract specific, hypothesis- and contrast-driven functional ROIs recruited when processing voice and noise separately (see Table 1 for an overview of the 26 ROIs extracted). We therefore focused on voice processing by contrasting vocal with non-vocal blocks—the latter blocks containing about 50% of environmental noise stimuli—and also performed the inverse contrast to focus on non-vocal, noise processing. These ROIs were then included in our functional and effective connectivity matrices as well as used as a mask in connectivity multi-voxel pattern analyses (fc-MVPA, see below).

**Table 1:**
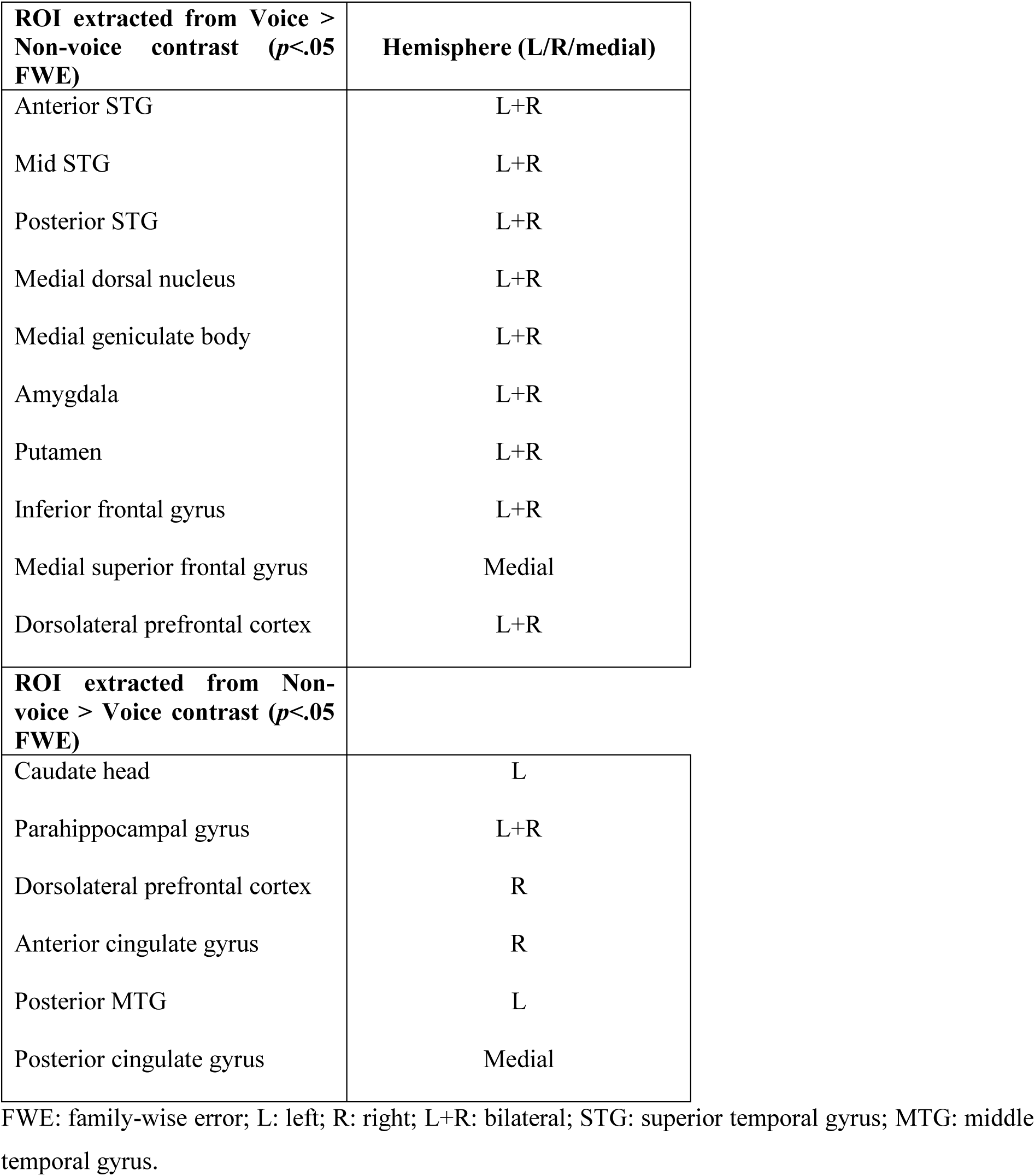
Regions of interest (ROIs) extracted for each whole-brain contrast of interest used in functional connectivity analyses.

| **ROI extracted from Voice > Non-voice contrast ($p < .05$ FWE)** | **Hemisphere (L/R/medial)** |
| --- | --- |
| Anterior STG | L+R |
| Mid STG | L+R |
| Posterior STG | L+R |
| Medial dorsal nucleus | L+R |
| Medial geniculate body | L+R |
| Amygdala | L+R |
| Putamen | L+R |
| Inferior frontal gyrus | L+R |
| Medial superior frontal gyrus | Medial |
| Dorsolateral prefrontal cortex | L+R |
| **ROI extracted from Non-voice > Voice contrast ($p < .05$ FWE)** |  |
| Caudate head | L |
| Parahippocampal gyrus | L+R |
| Dorsolateral prefrontal cortex | R |
| Anterior cingulate gyrus | R |
| Posterior MTG | L |
| Posterior cingulate gyrus | Medial |

FWE: family-wise error; L: left; R: right; L+R: bilateral; STG: superior temporal gyrus; MTG: middle temporal gyrus.

#### Whole-brain neuroimaging results

Voice processing—contrasting Voice to Non-Voice blocks—recruited large and distributed brain regions, notably along the temporal cortices bilaterally (Fig.1**a-c**). These expected enhanced temporal lobe activations included large portions of the anterior, mid and posterior superior temporal gyrus (STG) and the MTG, including most of the superior temporal sulcus as well (Fig.1**a,c**). These regions are known as the TVAs and our data reliably replicated the results of the princeps study^3^. We also found stronger activity in the bilateral prefrontal cortex, namely in the right inferior frontal gyrus (IFG), bilateral amygdala, putamen, medial dorsal nucleus and medial geniculate body of the thalamus (see Table 2). We extracted 19 ROIs from significantly enhanced voxels resulting from the Voice > Non-voice contrast (Table 1).

**Fig. 1:**
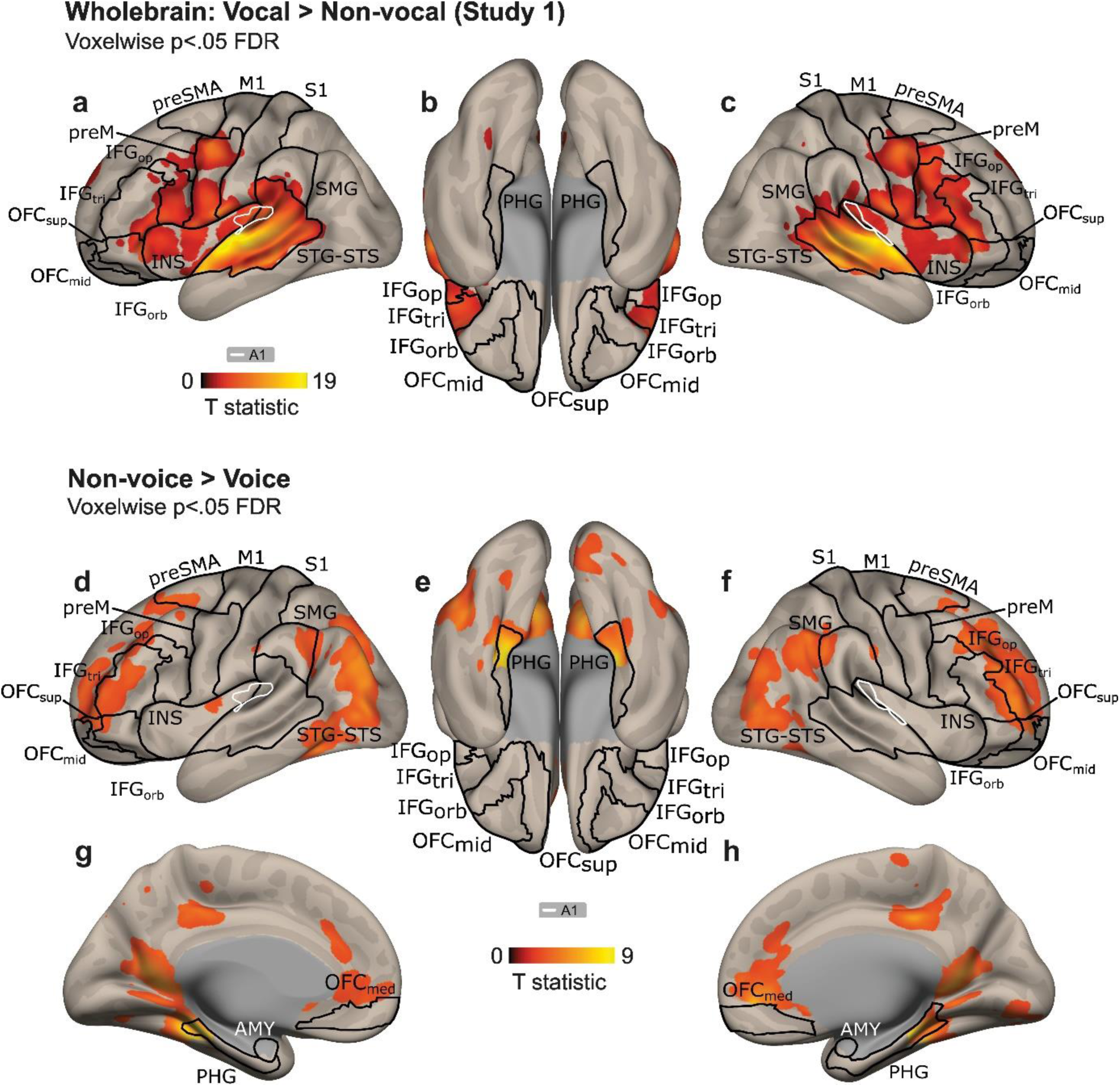
Enhanced brain activations specific to voice or noise processing. (**a-c**) Voice-specific processing in the bilateral middle and superior temporal cortex, planum temporale, inferior frontal gyrus and insula. (**d-h**) Non-voice-specific—including 50% of environmental noise stimuli—processing in the bilateral middle temporal gyrus, parahippocampal gyrus, inferior frontal gyrus, anterior & posterior cingulate cortices. All clusters and their coordinates are reported in Table 2. Statistically significant clusters are displayed on a normalized template at a threshold of *p*<.05, corrected for multiple comparisons at the voxel level (False Discovery Rate; FDR). The color bars illustrate the ‘t’ statistical values. A1: primary auditory cortex; IFG: inferior frontal gyrus; INS: insula; STG: superior temporal gyrus; STS: superior temporal sulcus; PHG: parahippocampal gyrus; AMY: amygdala; preSMA: pre-supplementary motor area; preM: premotor cortex; M1: primary motor cortex; S1: primary somatosensory cortex; SMG: supramarginal gyrus; OFC: orbitofrontal cortex. Suffixes: orb, pars orbitalis; tri, pars triangularis; op, pars opercularis; sup, superior; mid, middle; med, medial.

**Table 2:** Clusters and coordinates for Vocal > Non-vocal and Non-vocal > Vocal contrasts, Study 1, p<.05 FDR.

|  | Cluster size | T value | MNI X | MNI Y | MNI Z | Region | Hemisphere |
| --- | --- | --- | --- | --- | --- | --- | --- |
| **Vocal > Non-vocal** | 18128 | 20,49 | -62 | -18 | 2 | STG | L |
|  |  | 20,13 | 60 | -12 | -2 | STG | R |
|  |  | 19,78 | 60 | -2 | -4 | STG | R |
|  |  | 19,62 | -58 | -6 | -2 | STG | L |
|  |  | 16,56 | -60 | -30 | 6 | MTG | L |
|  | 605 | 9,39 | -50 | -6 | 46 | preM | L |
|  | 524 | 8,60 | 2 | 2 | 60 | preSMA | R |
|  | 156 | 6,41 | -24 | -62 | -20 | Cer VI | L |
|  | 703 | 5,30 | -4 | 56 | 28 | ACC | L |
|  | 556 | 4,75 | 4 | -58 | 32 | Precuneus | R |
|  | 84 | 4,28 | 22 | -60 | -18 | Cer VI | R |
|  | 58 | 3,46 | -6 | -14 | 42 | CCmid | L |
| **Non-vocal > Vocal** |  |  |  |  |  |  |  |
|  | 6159 | 9,39 | 26 | -30 | -18 | FFC | R |
|  |  | 8,93 | -26 | -38 | -16 | FFC | L |
|  |  | 8,75 | -32 | -32 | -14 | PHG | L |
|  |  | 7,38 | 14 | -50 | 10 | Calcarine | R |
|  | 3091 | 6,76 | 6 | 46 | -4 | OFCmed | R |
|  |  | 5,07 | 2 | 18 | -6 | Olfactory | R |
|  |  | 4,98 | -16 | 24 | 2 | Caudate | L |
|  |  | 4,61 | -34 | 58 | 4 | VLPFC | L |
|  |  | 4,54 | -12 | 16 | 12 | Caudate | L |
|  | 765 | 5,80 | 6 | -28 | 40 | CCmid | L |
|  |  | 4,46 | -6 | -38 | 42 | CCmid | L |
|  |  | 3,83 | 4 | -40 | 52 | Precuneus | L |
|  | 938 | 5,49 | 40 | -68 | 12 | MTG | R |
|  |  | 4,16 | 42 | -76 | 28 | LOCmid | R |
|  |  | 3,10 | 54 | -62 | 6 | MTG | R |
|  |  | 2,90 | 30 | -84 | 32 | LOCmid | R |
|  | 2381 | 5,36 | 38 | 28 | 38 | DLPFC | R |
|  |  | 5,35 | 32 | 52 | 2 | DLPFC | R |
|  |  | 5,30 | 40 | 50 | 10 | DLPFC | R |
|  |  | 5,29 | 30 | 36 | 34 | DLPFC | R |
|  | 857 | 4,71 | 52 | -48 | 50 | IPL | R |
|  |  | 4,45 | 56 | -46 | 40 | IPL | R |
|  |  | 4,21 | 44 | -48 | 34 | AG | R |
|  |  | 4,14 | 50 | -54 | 44 | IPL | R |
|  |  | 2,95 | 42 | -42 | 42 | SMG | R |
|  | 634 | 4,67 | -40 | 30 | 38 | DLPFC | L |
|  | 382 | 4,47 | -54 | -48 | 36 | IPL | L |
|  |  | 3,81 | -44 | -54 | 38 | IPL | L |
|  | 43 | 4,26 | 52 | -22 | 26 | SMG | R |
|  | 141 | 4,05 | 10 | -80 | -8 | Lingual | R |

STG: superior temporal gyrus; MTG: middle temporal gyrus; preM: premotor cortex; preSMA: pre- supplementary motor area; Cer: cerebellar lobule; ACC: anterior cingulate cortex; CC: cingulate cortex; FFC: fusiform cortex; PHG: parahippocampal gyrus; OFC: orbitofrontal cortex; VLPFC: ventrolateral prefrontal cortex; LOC: lateral occipital cortex; DLPFC: dorsolateral prefrontal cortex; IPL: inferior parietal lobule; AG: angular gyrus; SMG: supramarginal gyrus. Suffixes: mid, mid part.

The inverse contrast, i.e., Non-voice > Voice, yielded to enhanced BOLD signal in large portions of the bilateral prefrontal cortex, most notably in the IFG (Fig.1**d,f**), as well as in the anterior and posterior cingulate cortex (Fig.1**g,h**) and in the bilateral parahippocampal gyri (Fig.1**e**). Seven ROIs to be included in the connectivity analyses—and masked fc-MVPA analyses—were extracted from this second contrast (Table 1).

#### Connectivity results: Masked fc-MVPA

So far, we used mass-univariate statistical methods and here, we used multivariate analyses, taking the brain as a whole instead of running as many regressions as the number of existing voxels. Contrasting Voice to Non-voice trials led to enhanced multivariate connectivity in the left pMTG (posterior MTG), right IFGtri (IFG *pars triangularis*, Fig.2**a**) as well as in the posterior division of the bilateral parahippocampal gyrus (Fig.2**b**). These results allow for a clearer understanding of whole-brain data presented above, showing how connectivity patterns link lateral temporal, frontal and parahippocampal—medial temporal lobe—regions for voice processing.

**Fig. 2:**
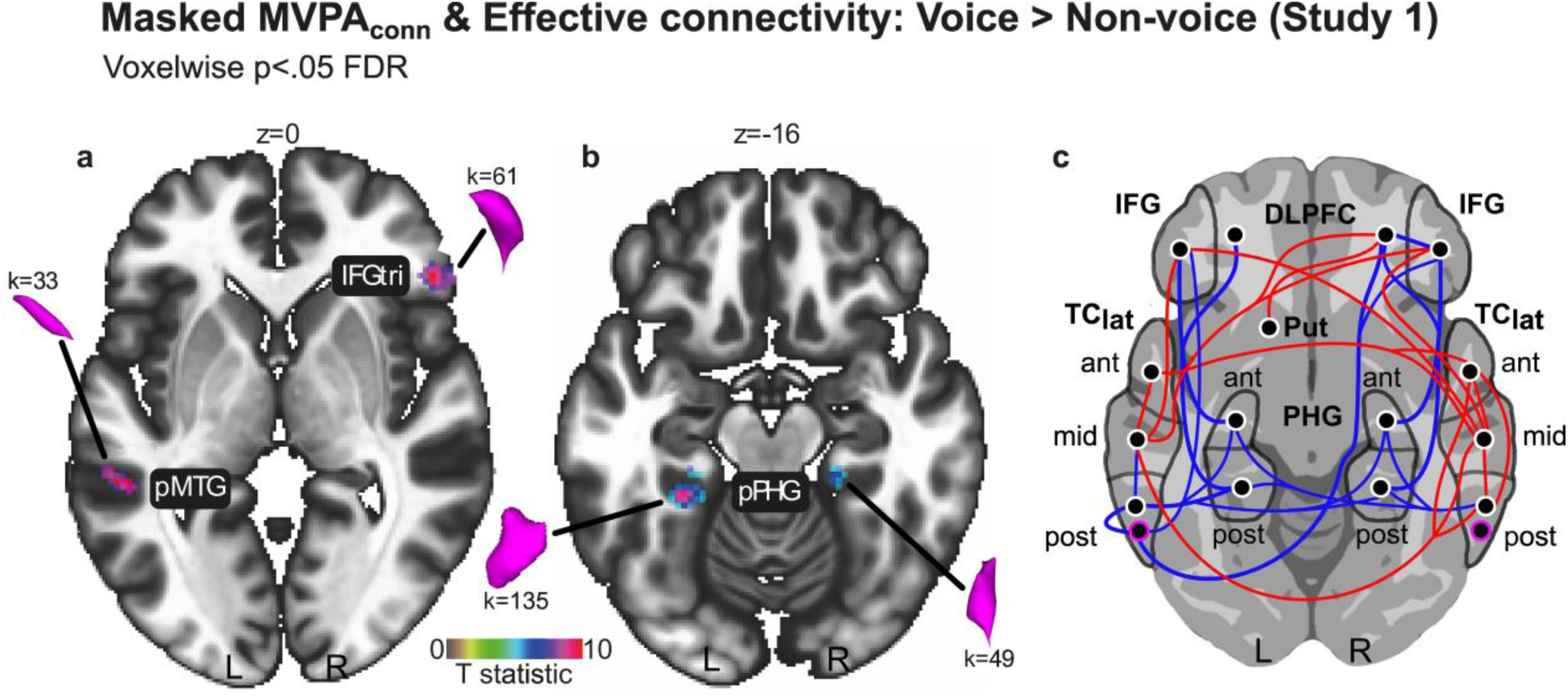
Effective connectivity underlying voice processing. (**a,b**) Connectivity multi-voxel pattern analysis for voice-specific processing. (**c**) Voice-specific, ROI-to-ROI effective connectivity—bivariate regression between ROIs, including both coupling (red lines) and anti-coupling (blue lines) between ROIs. The pink outlined circle in the posterior TC_lat_ regions indicates the posterior MTG (pMTG), the white outlined circle the posterior STG (pSTG). Effective connectivity was computed through generalized Psychophysiological Interaction analyses (gPPI). Statistically significant clusters are displayed on a normalized template at a threshold of *p*<.05, corrected for multiple comparisons at the voxel level (False Discovery Rate; FDR). The color bar illustrates the ‘t’ statistical values. IFGtri: inferior frontal gyrus, *pars triangularis*; pMTG: posterior middle temporal gyrus; pPHG: posterior parahippocampal gyrus; TC_lat_: lateral temporal cortex, including STG and STS regions; IFG: inferior frontal gyrus; DLPFC: dorsolateral prefrontal cortex; Put: putamen; PHG: parahippocampal gyrus; ant: anterior; post: posterior; mid: middle; L: left; R: right. K: number of voxels for a given cluster.

#### Seed-to-seed effective connectivity (generalized Psychophysiological Interactions using bivariate regression)

The second step of our connectivity analyses for Study1 was the comparison of seed-to-seed (or ROI-to-ROI) effective—and functional, see Fig.S1—connectivity for Voice against Non-voice blocks. We extracted a total of 26 ROIs on the basis of our two whole-brain contrasts of interest (19 for Voice > Non-voice and 7 for Non-voice > Voice; see Table 1). The general effective relations between these seed regions can be viewed in Fig.2**c** for the Voice > Non-voice contrast. Of note is that this contrast includes seed regions from both Voice > Non-voice and Non-voice > Voice whole-brain results (N=26 ROIs, as mentioned above). Coupling involved several connections in the lateral temporal cortex, mainly between the anterior, middle and posterior STG, bilaterally, and between the middle STG and the frontal cortex (IFG, dorsolateral prefrontal cortex: DLPFC), again bilaterally (Fig.2**c**). Anti-coupling involved a large network of ROIs, including again the IFG, DLPFC and more interestingly the anterior as well as posterior parahippocampal gyrus, the posterior STG and MTG, bilaterally (Fig.2**c**). ROI-to-ROI effective connectivity therefore highlights a coupled lateral temporal-frontal network while in parallel, an anti-coupled, medial temporal to lateral temporal cortex network was observed. See Table 3 for connectivity values.

**Table 3:**
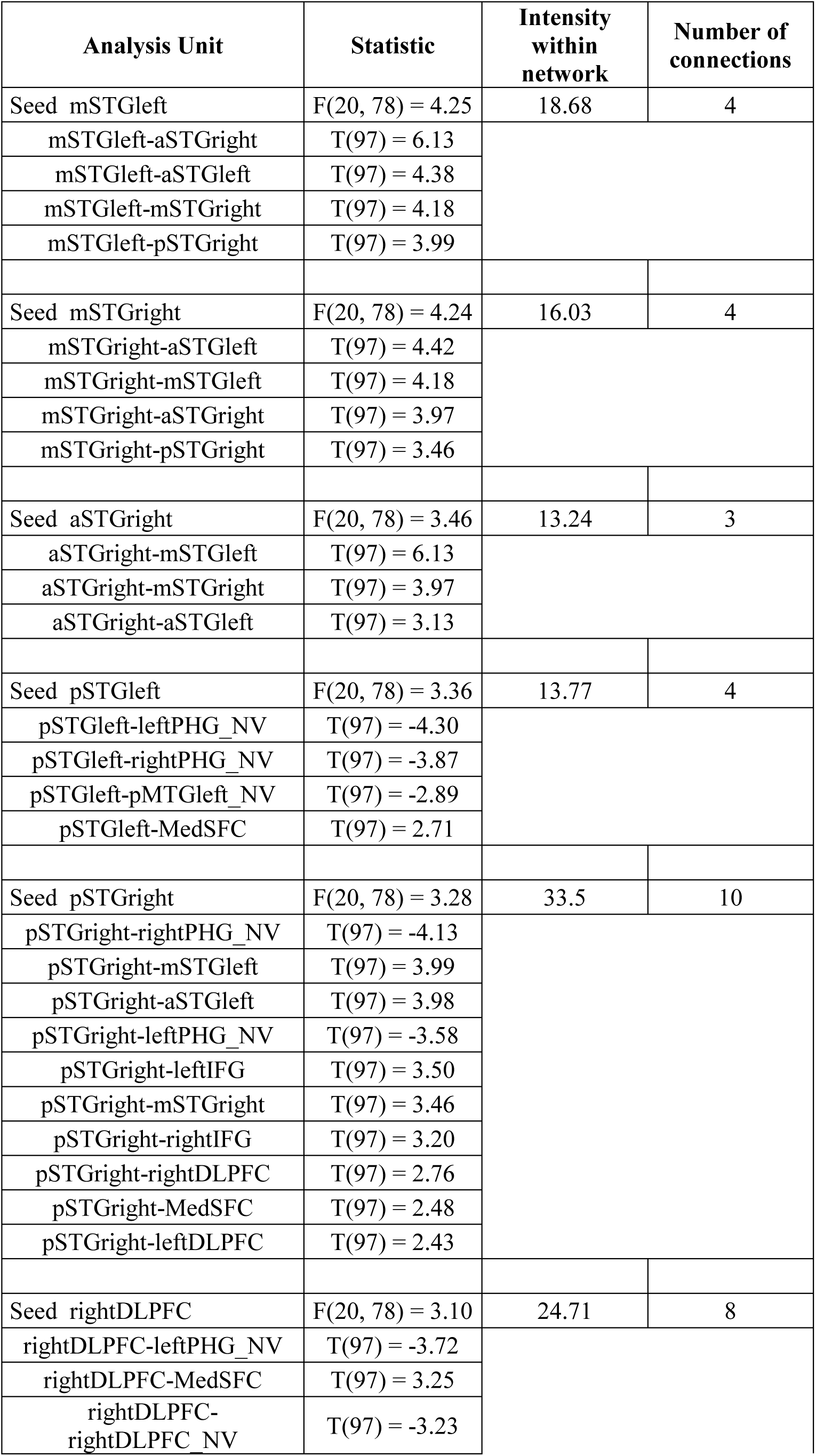
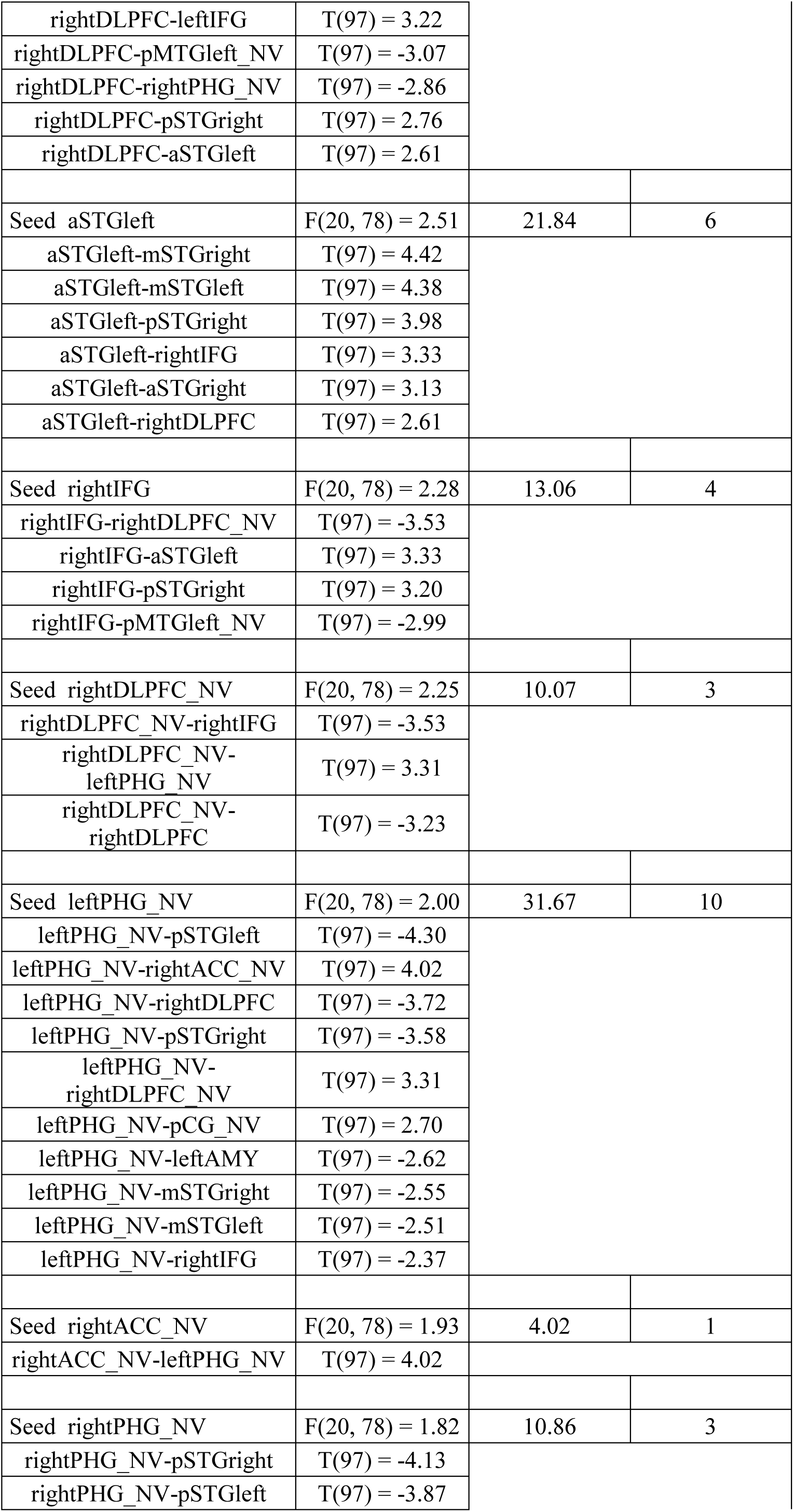
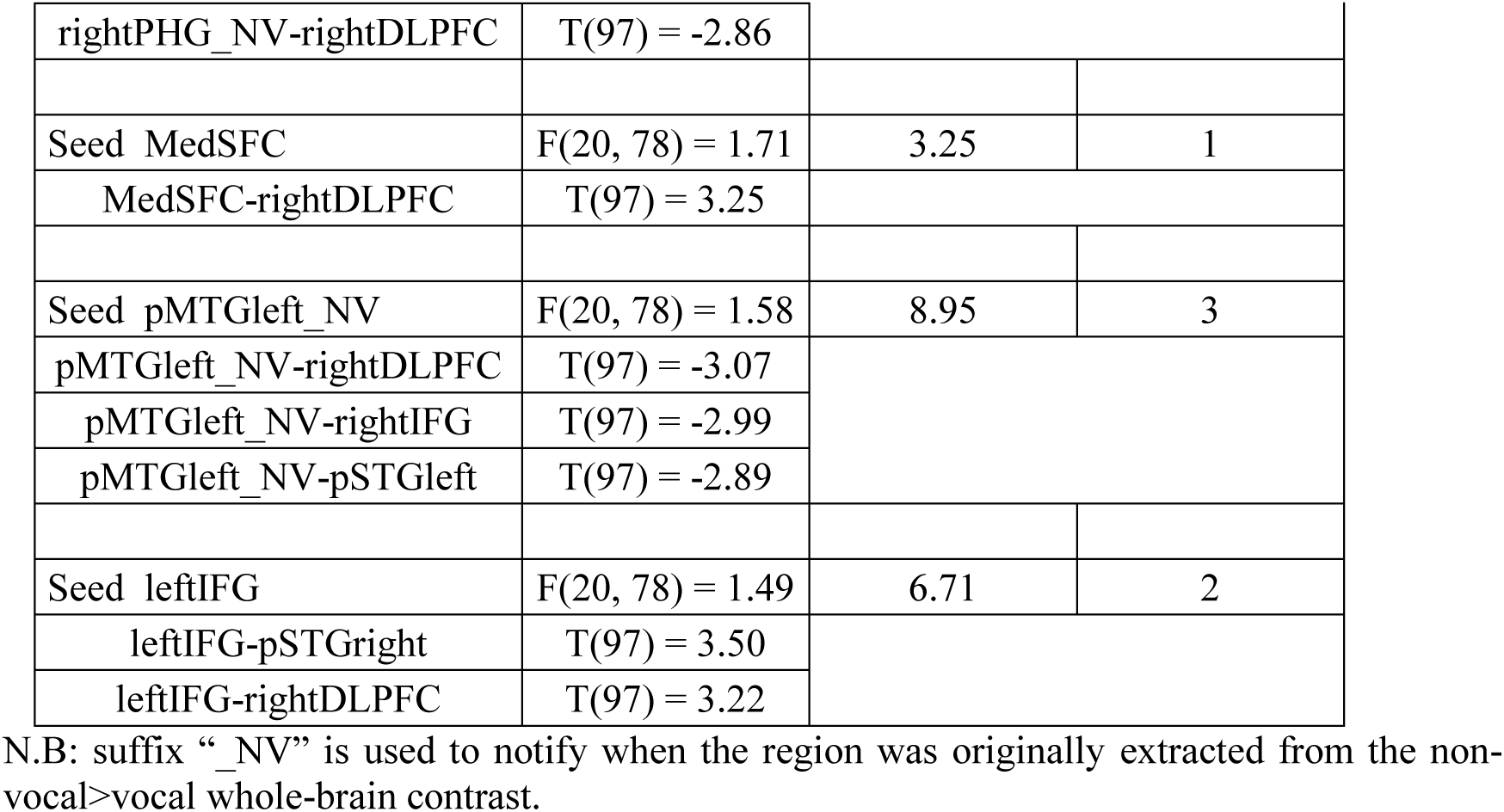
Matrix of connectivity values for seed-to-seed functional connectivity between all seed regions thresholded at *p*<.01 (analysis-wise false discovery rate, two-tailed).

Taken together, our effective connectivity results for Study 1 hence make it clear that voice processing not only involves the TVAs or lateral temporal cortex regions, but it also crucially relies on medial temporal connected areas such as the parahippocampal gyrus.

### Study 2 & 3: voice processing in noisy situations (behavioral study and neuroimaging)

Following the first study and the possibility that noise would be processed by the medial temporal lobe and the posterior MTG *during* voice processing, we raised the following question: are the MTL and pMTG working together to allow clear voice perception through—possibly parallel—noise reduction? Such question led to the necessity of directly manipulating background noise while concurrently perceiving voice signals, which was implemented in Study 2 as a behavioral pilot study and in Study 3 as an fMRI study. The paradigm of Study 3 was hence identical to the one used in Study 2—and included sparse-sampling MRI data acquisition to allow for silent periods while stimuli were presented and therefore improve general task sensitivity. In these studies, the participants had to specify whether the voice was easy or rather difficult to perceive in varying noise levels (Fig.3**b**). See the methods for more details, Fig.3**a** below for task design and Fig.3**b** for conditions as well as stimulus characteristics and manipulation.

**Fig. 3:**
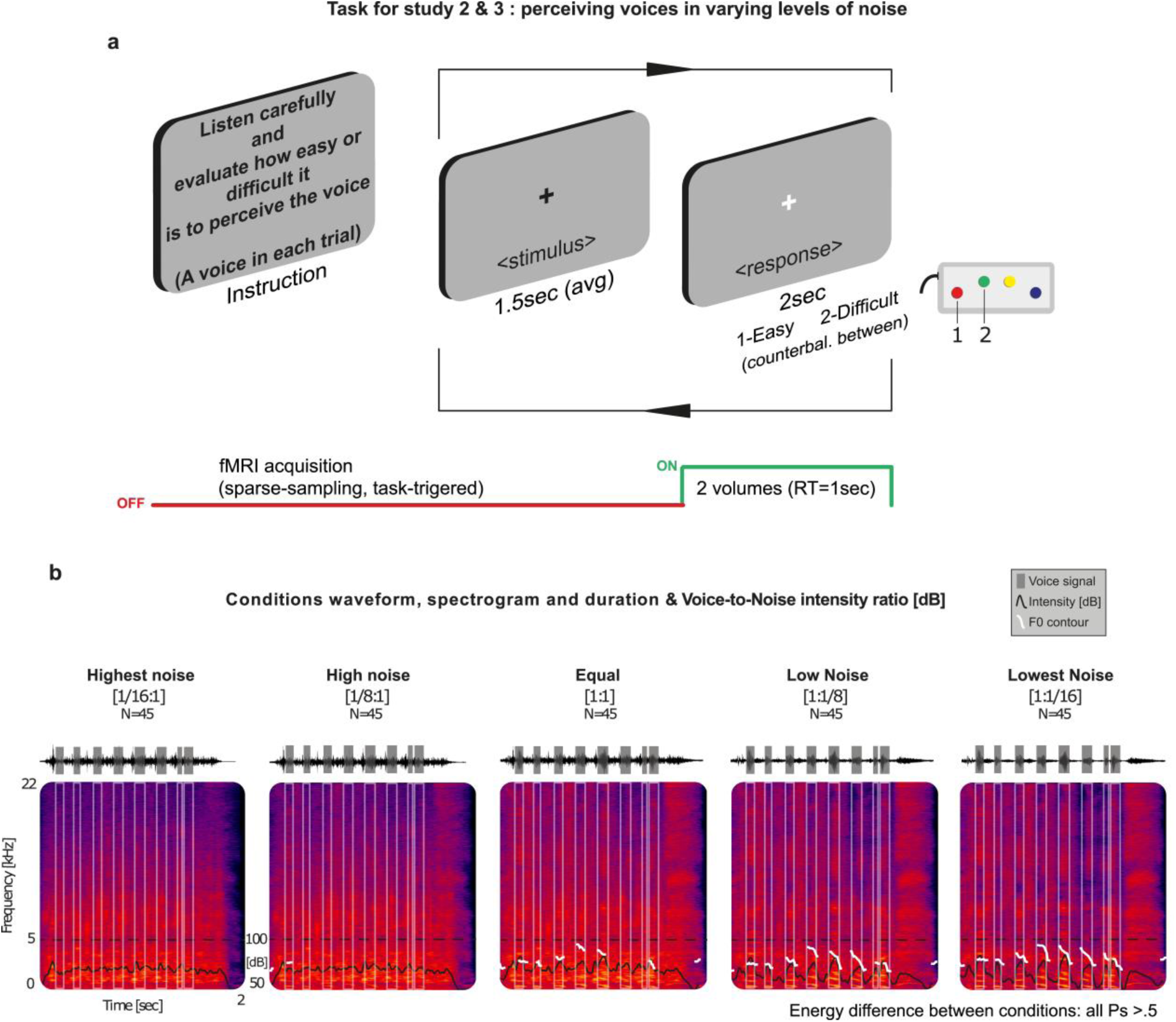
Task design and stimulus description for Study 2 and 3. (**a**) Task design for Study 2 & 3, illustrating task instruction and one trial. Stimulus was presented for 1.5sec in average and response time was fixed to 2sec, with a 2 Alternate-Forced Choice response (buttons counter-balanced between participants). Participants had to be as accurate as possible, but not as fast as possible since precision was the goal here. Study 2 was behavioral only while Study 3 included MRI scanning in a script-triggered, sparse-sampling fashion to avoid additional sources of noise during stimulus perception. Two full volumes of 1sec each were therefore acquired during the response slide, for 2sec in total. (**b**) Stimuli consisted of a combination of voice and environmental noise, carefully selected from the Non-voice stimuli of the database of Study 1^1^. There were five conditions (Highest noise, High noise, Equal, Low noise, Lowest noise), each including 45 trials (total N_trials_=225) and a specific ratio of voice-to-noise intensity (difference between conditions concerning energy showed no significant effect with all Ps > .5, see Methods). fMRI: functional magnetic resonance imaging; RT: MRI repetition time; dB: decibel; F0: voice fundamental frequency; kHz: kilohertz (1Hz*1000=1kHz).

#### Probability of perceiving voice signals in varying levels of noise (behavior for Study 2 & 3)

Behavioral results of both studies 2 & 3 illustrate the probability of perceiving a voice in a noisy background and were computed using mixed effects, logistic regression analyses (see Methods for details). The dependent variable was the response of the participant, the fixed effect was the level of noise and random effects included the identity of the participant, its gender and age as well as the identity and order of each stimulus presented. Response probability shows a clear degradation of the perception of voice signals in noise when noise level is High to Highest (voice-to-noise dB ratio=1/8:1 & 1/16:1, respectively) in both independent samples (Fig.4**a**, left panel: Study 2, N=18; right panel: Study 3, N=20). Individual data values and violin distribution curves for reaction times data are reported in Fig.S2.

**Fig. 4:**
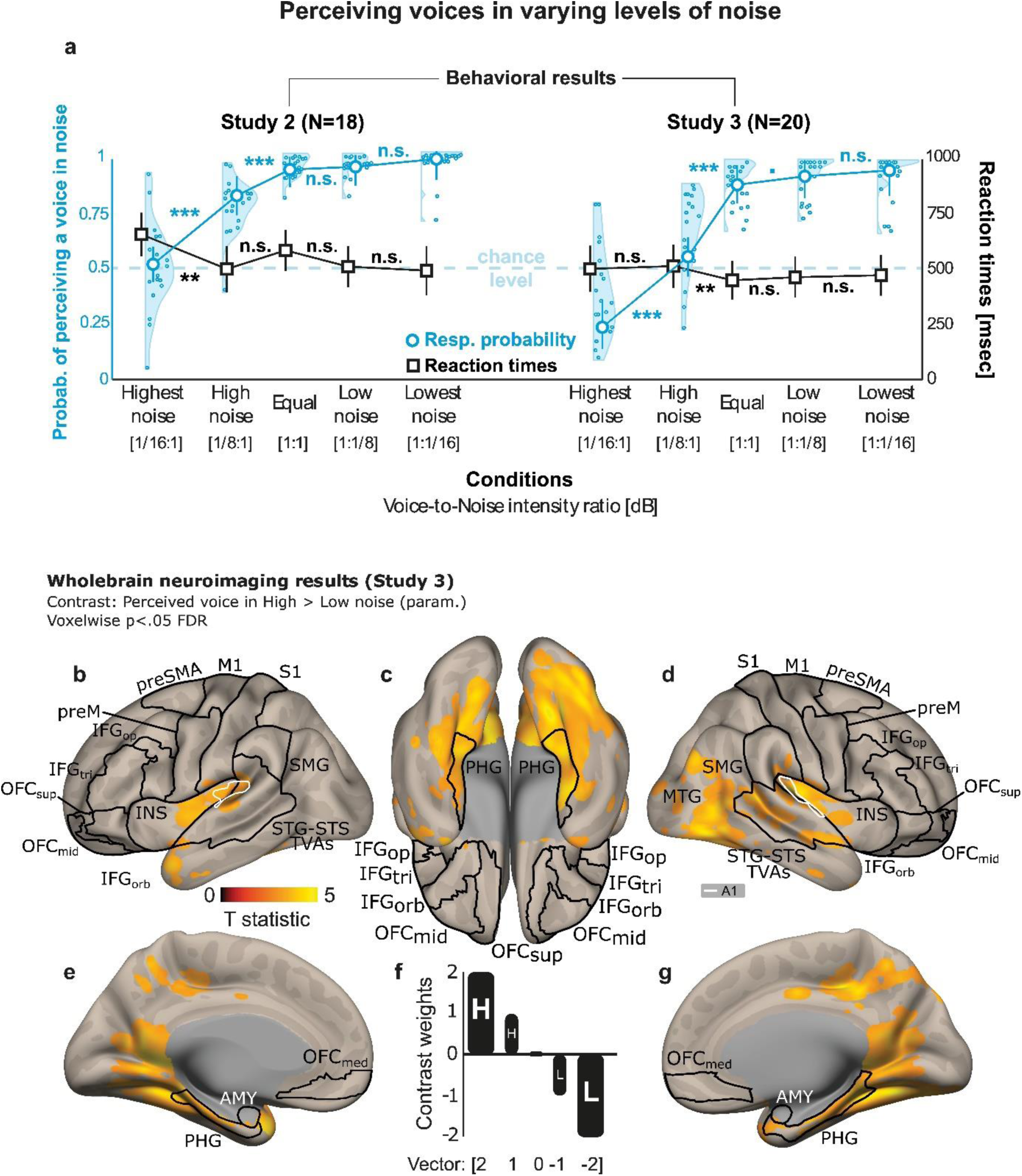
Perception of voice signals in varying levels of noise, behavior and neuroimaging (Study 2 & 3). (**a**) Behavioral results of both studies illustrating the probability of perceiving a voice in a noisy background (logistic regression analysis, in blue, circles indicate individual values, half-violin curves the distribution of the data) as well as reaction times (in black, squares, individual values and half-violin curves in Fig.S2), *y* axis. Response probability shows a clear degradation of the perception of voice in noise when noise level is High to Highest in both independent samples (left panel: Study 2, N=18; right panel: Study 3, N=20). X axis: conditions with voice-to-noise ratio, in decibel (dB). Error bars represent the standard error of the mean. Chance level is 50%. n.s.: not significant; **·**: statistical tendency (.05<p<.1); \*\*\**p*<.001; \*\**p*<.01. Probab.: probability; Resp.: response. (**b-e, g**) Whole-brain neuroimaging results of voice perception in noisy situations highlighting posterior lateral temporal cortex (**d,e,g**) and bilateral parahippocampal activity (**c**) as specified by (**f**) the parametric contrast: [2(Highest noise) 1(High noise) 0(Equal) -1(Low noise) -2(Lowest noise)]. All clusters and their coordinates are reported in Table 4. Statistically significant clusters are displayed on a normalized template at a threshold of *p*<.05, corrected for multiple comparisons at the voxel level (False Discovery Rate; FDR). The color bars illustrate the ‘t’ statistical values. A1: primary auditory cortex; IFG: inferior frontal gyrus; INS: insula; MTG: middle temporal gyrus; STG: superior temporal gyrus; STS: superior temporal sulcus; PHG: parahippocampal gyrus; AMY: amygdala; preSMA: pre-supplementary motor area; preM: premotor cortex; M1: primary motor cortex; S1: primary somatosensory cortex; SMG: supramarginal gyrus; OFC: orbitofrontal cortex; TVAs: temporal voice areas. Suffixes: orb, pars orbitalis; tri, pars triangularis; op, pars opercularis; sup, superior; mid, middle; med, medial.

In Study 2, we observed a significant impact of noise levels on the probability of perceiving a voice signal (χ^2^(4)=283.93, *p*<.001). Variance explained by this model was 31.68% for the fixed factor only (R^2^m) and 61.75% when including both fixed and random effects (R^2^c). Using a confidence interval of 97.5% and a Bonferroni correction for multiple comparisons, we computed paired contrasts of interest between each level of noise (Fig.4**a**, right panel). Results show no difference between Lowest and Low noise (χ^2^(1)=0.06, *p*=.81) and between Low noise and Equal voice and noise levels (χ^2^(1)=0.83, *p*=.36), significant differences between Equal and High noise (χ^2^(1)=47.81, *p*<.001) as well as between High and Highest noise levels (χ^2^(1)=52.12, *p*<.001), interestingly. Using a parametric contrast including all conditions and targeting voice perception in noisy as opposed to less noisy situations—with a stronger difference between Highest vs. High and Lowest vs. Low values—led to a significant effect (χ^2^(1)=208.28, *p*<.001). The vector of this contrast was the following: [2(Highest noise) 1(High noise) 0(Equal) -1(Low noise) -2(Lowest noise)]. Therefore, the probability of reporting the voice was ‘easy’ as opposed to ‘difficult’ to perceive was modulated by the level of background noise, especially when noise was Highest and High compared to when it was Low or Lowest.

Concerning reaction times—even though of no particular interest here, we observed a significant effect of noise level on response probability (χ^2^(4)=16.03, *p*<.01), triggered by a difference between the Highest and the High noise conditions (χ^2^(1)=8.45, *p*<.01). No other contrast reached significance (Lowest > Low noise: χ^2^(1)=0.04, *p*=.84; Low noise > Equal: χ^2^(1)=0.85, *p*=.36; Equal > High noise: χ^2^(1)=0.57, *p*=.45). We again used the same parametric contrast as above, including all conditions and targeting voice perception in noisy as opposed to less noisy situations—with a stronger difference between Highest vs. High and Lowest vs. Low values—and it led to a significant effect (χ^2^(1)=8.45, *p*<.01). This result highlights slower reaction times between high noise situations, which seems to indicate higher task difficulty as well. Again, the confidence interval was 97.5% and a Bonferroni correction was used. Variance explained by this model was 0.39% for the fixed factor only (R^2^m) and 6.22% when including both fixed and random effects (R^2^c).

In fMRI Study 3, we observed—as in behavioral Study 2—a significant impact of noise levels on the probability of perceiving a voice signal (χ^2^(4)=375.65, *p*<.001). Variance explained by this model was 24.08% for the fixed factor only (R^2^m) and 71.43% when including both fixed and random effects (R^2^c). Using a confidence interval of 97.5% and a Bonferroni correction for multiple comparisons, we computed paired contrasts of interest between each level of noise (Fig.4**b**, right panel). Results show no difference between Lowest and Low noise (χ^2^(1)=0.57, *p*=.45), a tendency between Low noise and Equal voice and noise levels (χ^2^(1)=3.69, *p*=.055) and significant differences between Equal and High noise (χ^2^(1)=67.44, *p*<.001) as well as between High and Highest noise levels (χ^2^(1)=77.67, *p*<.001). Using the same parametric contrast as in Study 2, we observed again a significant effect (χ^2^(1)=314.01, *p*<.001). Therefore, we can again say here that the probability of reporting the voice was ‘easy’ as opposed to ‘difficult’ to perceive was modulated by the level of background noise, especially when noise was High and Highest compared to when it was Low or Lowest.

Concerning reaction times—and even though they were still of no interest in Study 3, we observed a significant effect of noise level on response probability (χ^2^(4)=18.59, *p*<.001), triggered by a difference between an Equal ratio of voice-to-noise and the High noise condition (χ^2^(1)=8.59, *p*<.01). No other contrast reached significance (Lowest > Low noise: χ^2^(1)=0.02, *p*=.89; Low noise > Equal: χ^2^(1)=0.015, *p*=.90; High > Highest noise: χ^2^(1)=0.002, *p*=.97). We again used the same parametric contrast as above, and it led to a significant effect (χ^2^(1)=12.95, *p*<.001). This result highlights slower reaction times for High—but not Highest—noise situations, which again seems to indicate higher task difficulty as well. Again, the confidence interval was 97.5% and a Bonferroni correction was used. Variance explained by this model was 0.42% for the fixed factor only (R^2^m) and 15.24% when including both fixed and random effects (R^2^c).

#### Whole-brain neuroimaging results (Study 3)

Neuroimaging results were obtained using the sample of Study 3, since Study 2 was only a behavioral study. Here, we were specifically interested in using High- as opposed to Low-noise situations. Therefore, and to be consistent with behavioral analyses, we used the same parametric contrast as above. It included all conditions and targeted voice perception in noisy as opposed to less noisy situations— with a stronger difference between Highest vs. High and Lowest vs. Low trials. The vector of this contrast was identical to the one used for behavior (Fig.4**f**), namely: [2(Highest noise) 1(High noise) 0(Equal) -1(Low noise) -2(Lowest noise)]. Trials in which the participants indicated they easily perceived the voice signals were used, while those ‘difficult’ trials were concatenated into a single column of no-interest, since we wanted to understand ‘successful’ voice processing when noise is present. The data emphasize a strong role of the right posterior MTG (Fig.4**d**), anterior and posterior bilateral PHG (Fig.4**c,e,g**), right posterior, mid and anterior STS (Fig.4**d**) and bilateral insula (Fig.4**b,d**). All above-threshold clusters are reported in Table 4. The inverse contrast yielded to vast prefrontal, orbitofrontal and inferior activity, but posterior MTG or any of the PHGs did not show any above-threshold enhanced activity (Fig.S3). Among the regions of the main contrast of interest, it is important to highlight that many of them overlap with both Voice-specific and Non-voice-specific brain areas observed in Study 1 (see Fig.S4 and Fig.S5, respectively), therefore adding more weight to the assumption of their parallel role for processing voice in noise. The F contrast including all trials (‘easy’ and ‘difficult’ voice perception as assessed by our participants’ responses) is reported in Fig.S6, while the F contrast of solely the ‘easy’ perception trials is reported in Fig.S7.

**Table 4:** Clusters and coordinates for Voices perceived in High compared to Low noise situations, Study 3, p<.05 FDR.

| Cluster size | T value | MNI X | MNI Y | MNI Z | Region | Hemisphere |
| --- | --- | --- | --- | --- | --- | --- |
| 21795 | 4,91 | 26 | -60 | -10 | Lingual | R |
|  | 4,80 | 50 | -66 | -8 | ITG | R |
|  | 4,69 | -12 | -46 | 4 | Calcarine | L |
|  | 4,58 | -16 | -56 | 6 | Calcarine | L |
|  | 4,58 | 42 | -72 | -10 | FFC | R |
| 957 | 4,82 | 14 | -28 | 40 | CCmid | R |
|  | 3,96 | 4 | -56 | 52 | Precuneus | R |
|  | 3,95 | -16 | -36 | -14 | PHG | L |
|  | 3,72 | 14 | -84 | 46 | Cuneus | R |
|  | 3,65 | -10 | -30 | 40 | CCmid | L |
|  | 3,57 | 22 | -34 | -12 | PHG | R |
|  | 3,45 | -4 | -54 | 44 | Precuneus | L |
| 153 | 4,24 | 40 | -78 | 34 | LOCmid | R |
| 469 | 4,03 | -46 | -10 | -2 | STG | L |
|  | 3,55 | -36 | -4 | 8 | Insula | L |
|  | 3,49 | -40 | -20 | -2 | STG | L |
| 39 | 3,69 | 4 | -6 | 38 | CCmid | R |
| 34 | 3,16 | -46 | 8 | -20 | STG | L |
| 33 | 3,12 | 26 | -94 | 16 | LOCsup | R |

STG: superior temporal gyrus; CC: cingulate cortex; FFC: fusiform cortex; PHG: parahippocampal gyrus; LOC: lateral occipital cortex; ITG: inferior temporal gyrus. Suffixes: mid, mid part; sup, superior part.

#### Connectivity results (Study 3): Masked fc-MVPA and seed-to-voxel functional and effective connectivity (generalized Psychophysiological Interactions)

In order to grasp the full picture of voice processing in noisy situations, we then turned to undirected— this section, Fig.5—and directed connectivity analyses—dynamic causal modeling, next section (Fig.6) using the same integrative contrast as before ([2(Highest noise) 1(High noise) 0(Equal) -1(Low noise) - 2(Lowest noise)]). We started by computing connectivity MVPA and this multivariate method yielded to a single region, namely the right posterior MTG (Fig.5**a**). This region of course greatly overlaps with the one observed in whole-brain results of this study (Fig.4**d**). Taking this right pMTG as seed for further seed-to-voxel effective connectivity analyses using bivariate regressions, results show a direct negative relation with the left PHG, more specifically its posterior division (Fig.5**b**). To close the circle, we then used this right posterior PHG as seed. We also included the left pPHG observed in Study 1 (specific to Non-voice) to better disentangle a potential dissociation between hemispheres in the PHG. Both functional—using bivariate correlation—and effective—using bivariate regression—connectivity analyses were performed here. The left pPHG showed functional anti-coupling back with the right pMTG (Fig.5**c**) while the right pPHG showed effective, direct anti-coupling with the right planum temporale (Fig.5**d**). Taken together with the results of Study 1 as well as those mentioned above of Study 2 and of this study, these results give even more strength to the concept of a medial temporal network working in concert with the TVA and voice-processing areas for an improved voice perception in noisy situations.

**Fig. 5:**
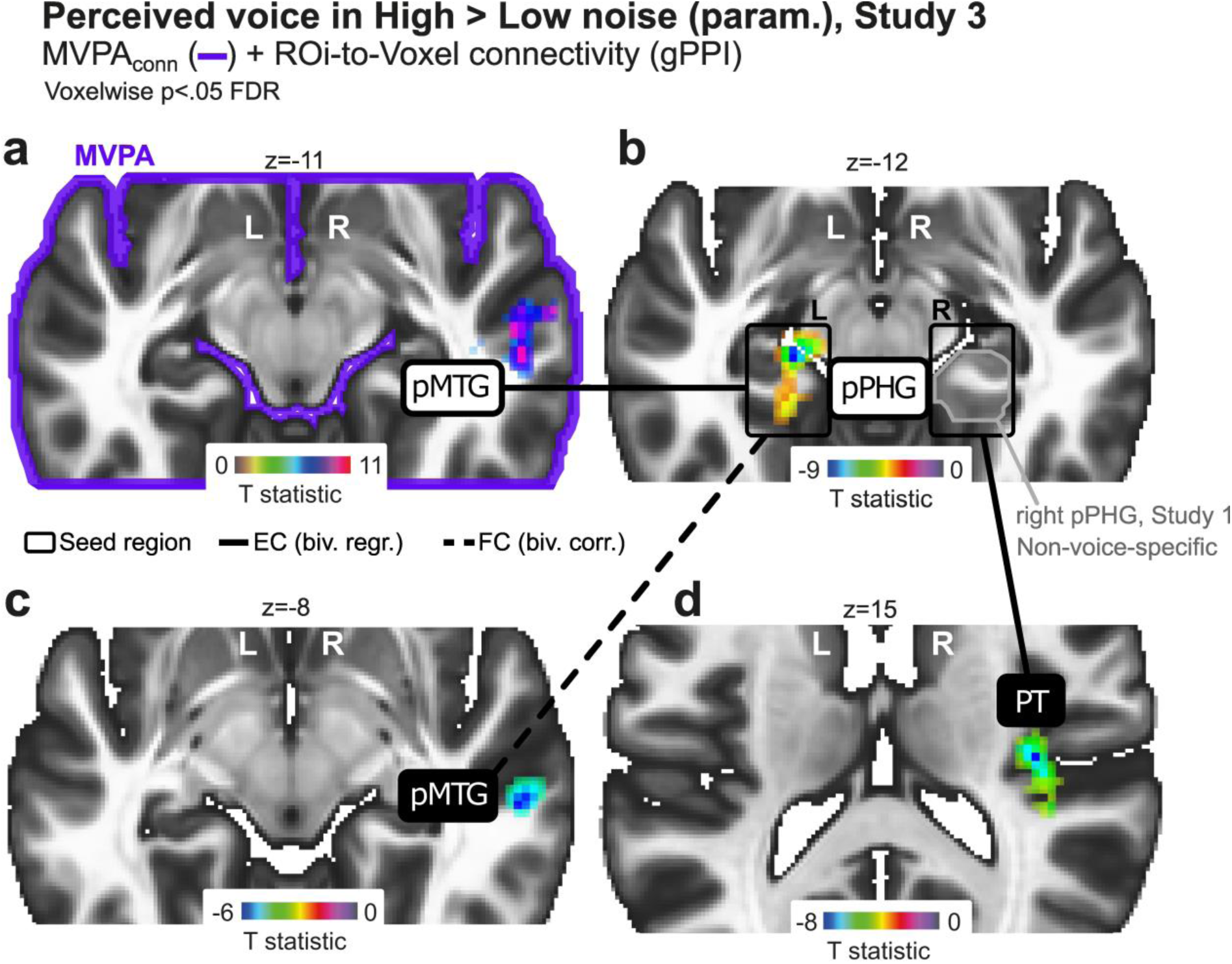
Effective and functional connectivity underlying voice processing in noise. (**a**) Connectivity multi-voxel pattern analysis (MVPA) resulting in right posterior middle temporal gyrus (pMTG) used as seed for (**b**) effective connectivity analysis leading to a direct anti-coupling with the left posterior parahippocampal gyrus (pPHG), itself functionally anti-coupled back with the right pMTG (**c**). (**d**) When taking the right pPHG—observed in study 1 for the Non-voice-specific contrast, as seed, we found direct anti-coupling with the right planum temporale (PT). Effective connectivity used bivariate regression while functional connectivity used bivariate correlation, both computed through generalized Psychophysiological Interaction analyses (gPPI). Statistically significant clusters are displayed on a normalized template at a threshold of *p*<.05, corrected for multiple comparisons at the voxel level (False Discovery Rate; FDR). The color bars illustrate the ‘t’ statistical values. L: left; R: right.

**Fig. 6:**
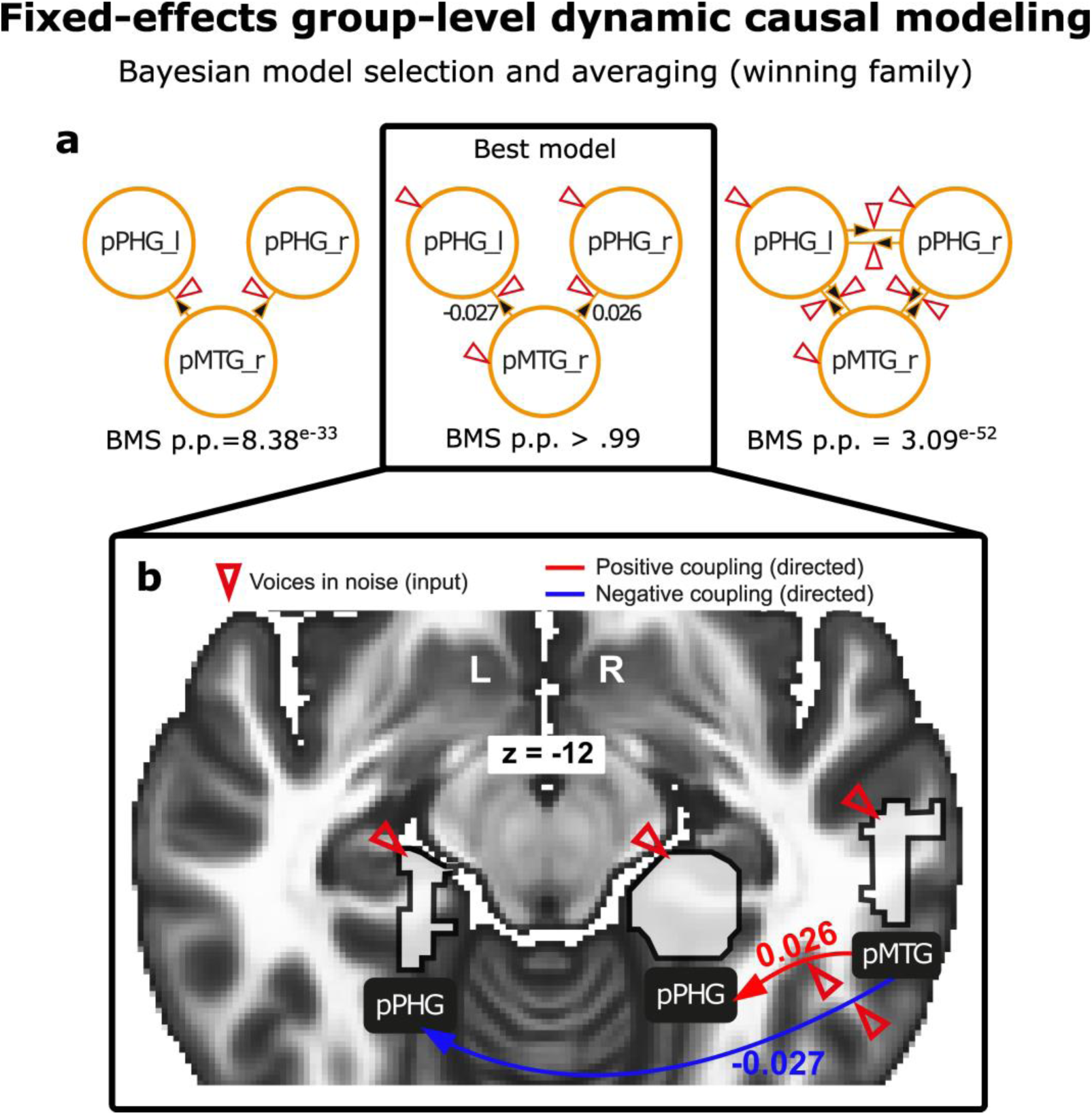
Effective, directed fixed-effect group-level connectivity underlying voice processing in noise using dynamic causal modelling (DCM). DCM analysis was used on nine models and three families, all containing the right posterior middle temporal gyrus (pMTG) and the bilateral posterior parahippocampal gyrus (pPHG). Families and models are detailed in Fig.S8. (**a**) The input was represented by High and Highest noise conditions (red triangle), and (**b**) direct coupling was observed from the right pMTG to the right pPHG (red arrow) while direct anti-coupling was observed from the right pMTG to the left pPHG (blue arrow). Bayesian model averaging and Bayesian model selection criteria both surpassed a posterior probability of p>.99. L: left; R: right.

#### Dynamic causal modelling, fixed effects (Study 3)

We computed one last analysis for this study, with the aim of constraining our data even more to understand how the relations between the lateral and medial temporal cortex would constitute a biological mechanism recruited for voice perception in noise. To do so, we carefully designed nine credible models—in addition to one for defining intrinsic connectivity—decomposed into three families based on their given input (see Fig.S8). The winning family—involving High and Highest noise conditions as input to the right pMTG and bilateral pPHG—contained the winning model with condition input to the right pMTG, the bilateral pPHG as well as input to the directed effective relation from the right pMTG to the bilateral pPHG (Fig.6**a**,**b**). Because we were interested in within-participant rather than between-participant differences, we computed a fixed-effect analysis type. For deciding the winning family, the Bayesian model averaging posterior probability was above pp>.99, same goes for Bayesian model selection allowing model 8 to be the selected as the best model (pp>.99, see Fig.6**b** and Fig.S8 for more details). The winning model therefore had inputs of High and Highest noise conditions on all three regions of interest as well as on the positive directed link from the right pMTG to the right pPHG and on the negative directed link from the right pMTG to the left pPHG (Fig.6**a**). Finally, it is important to note that in this analysis, the ROIs were the actual clusters found in the previous results sections mentioned above, not a sphere or a square average based on coordinates (see Methods).

## Discussion

The present studies aimed at a better understanding of voice processing at the neural level with the general goal of shedding new light on auditory and language pathways used by humans in social interactions and in everyday life communication. We used a combination of tasks involving the processing of voice and noise3 separately (Study 1)—as well as voice *in* noise (Study 2 & 3), by modulating the background environment of auditory scenes. In Study 1, we observed two distinct neural networks underlying the processing of vocal and non-vocal auditory material—the anterior, mid, posterior superior temporal cortex and the IFG for voice processing; the posterior STC and the bilateral parahippocampal gyrus for the processing of non-vocal stimuli. The latter represents a novel anti-coupled subnetwork, originating from the posterior part of the parahippocampal gyrus (bilaterally) to the most posterior part of the bilateral temporal lobe (middle and superior portions). Are these brain networks working hand in hand to allow clear voice perception in noise? Is this latter network playing any role in the boosting of voice processing in noisy situations? Is it responsible for noise reduction?

To clarify these aspects—that could not be addressed by the design of Study 1, a second task was programmed and performed on independent samples both behaviorally and in an fMRI scanner (Study 2 & 3, respectively; sparse-sampling fMRI in Study 3), involving voice perception in varying levels of realistic background noise. The results revealed expected behavioral effects—the more background noise, the more difficult it becomes for the participants to distinguish vocal signals, in addition to neural effects greatly overlapping with those observed in Study 1: voice perception in high levels of environmental noise triggered enhanced brain activity in the posterior temporal lobe as well as in the bilateral parahippocampal gyrus—named throughout this section lateral (TC_lat_) and medial (TC_med_) temporal cortex, respectively. The commonalities between our fMRI studies led us to the articulation of our findings as a specific model of voice processing in noisy situations, which is summarized in Fig.7 below. We will iterate this model throughout this section and discuss in further detail the implications of our results for social neuroscience and mention future perspectives to consider.

**Fig. 7:**
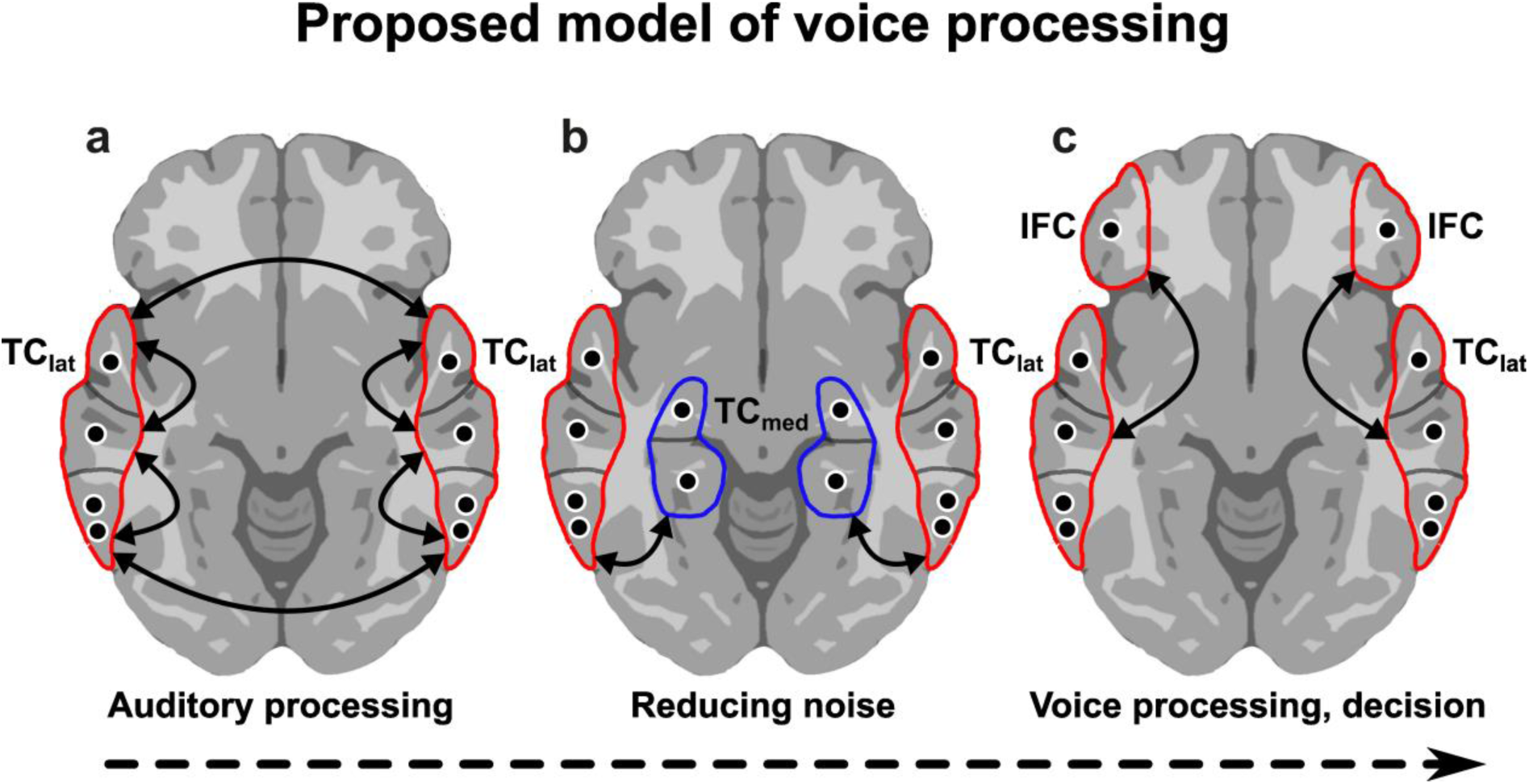
Proposed model of voice processing in noisy situations. Based on the results obtained in the present set of studies (Study 1-3), we postulate an updated model of voice processing, more specifically addressing the processing and possible parallel diminution of background noise. Although fMRI does not have a temporal resolution dynamic enough to focus on the time-course of voice processing in noise, we present three steps, happening in a supposed combination of serial and parallel steps: (**a**) First, auditory processing happens in the auditory cortex, namely the lateral temporal cortex (TC_lat_), both in a bilateral and ipsilateral fashion through coupling (red outline). (**b**) Second, background noise is reduced thanks to the anti-coupling (blue outline) between voice-specific TC_lat_ regions and noise-specific medial temporal cortex (TC_med_) regions. (**c**) Lastly, voice processing ‘free of noise’—or at least with the main sources of noise diminished enough to allow for clear voice perception—resumes and/or continues in parallel and decision processes can be triggered thanks to a communication between TC_lat_ and the inferior frontal cortex (IFC). This step may well occur partially in parallel to step 2, and step 2 may well occur partially in parallel to step 1.

Among the evolutionary changes of modern humans, verbal communication was—and still is—of the highest importance. The vector of human verbal communication being the vocal signal, the study of the human voice appears therefore to be a crucial research topic and this argument represents the main motivation behind the present body of work. In addition to vocal verbal communication, suprasegmental aspects (i.e., pitch, voice quality, intonation) of the human voice contain information not only about the emotional state of the speaker^34,35^, but also about its identity^36^, age and gender^37^, and even personality^38,39^. In parallel to these highly informative behavioral aspects, neuroimaging highlighted the role of the temporal lobe (i.e., TVAs)^3^ in voice processing and our results are in line with these *princeps* findings^34^. More specifically, three sub-clusters of the TVAs were observed in our data, namely the posterior, mid and anterior regions, and these TVAs subparts were positively correlated (Fig.2**c**), a result that was already reported previously^4^. However, our data revealed that the most posterior part of the STC correlated with the anterior and mid parts of the STC in a weaker manner, especially the left posterior STG. Since the posterior auditory cortex or posterior STC was shown to gate novel sounds to consciousness^40^, to underlie the recognition of environmental sounds^19^ and to process voice as part of the TVAs, why would this region be less connected to the rest of the temporal lobe during voice processing? More importantly, the bilateral posterior STC—including the MTG—was also part of an anti-coupled network together with the parahippocampal gyrus, bilaterally. The parahippocampal gyrus was reported to mediate the indirect processing of vocal threat^21^, as well as the perception and identification of environmental sounds^41^, but its role in general voice perception has, to the best of our knowledge, neither been revealed nor explored. Therefore, we must understand the role of the parahippocampal gyrus for voice perception—since we know its involvement in noise perception, most crucially in relation to the posterior temporal cortex. A specific investigation of voice perception in noise therefore had to be assessed—since we think noise could be reduced or even filtered out thanks to this anti-coupled TC_lat_-TC_med_ network, and such aspect was not manipulated at all in Study 1.

The assumption that a network of regions including the parahippocampal gyrus (TC_med_) and the posterior STC and MTG (TC_lat_) might play a role in voice processing in noise should therefore yield to enhanced activity and/or connectivity when voices are processed and well perceived in high vs low noise situations. This is exactly what we observed in Study 3 both for wholebrain and connectivity analyses (functional, effective—both for directed and undirected measures). Our results together with existing literature on the perception of voice^3,4,9,14-16,20,21,27,36^, noise^17,19,23,25,26,41^ and voice-in-noise^18,33,42-45^ processing led us to drafting a three-step neural model of voice processing in noise (Fig.7). As mentioned above, we refer to the bilateral TVAs and especially the STC and MTG as the ‘lateral temporal cortex’ (TC_lat_) while we regroup the subregions of the parahippocampal gyrus as the ‘medial temporal cortex’ (TC_med_). Inferior frontal cortex subregions (*pars orbitalis*, *opercularis* and *triangularis*) are also regrouped under the general term ‘IFC’. Three steps would unfold to allow for efficient and accurate voice processing in noise and potentially for any type of social interactions involving speech. These steps include: i) the processing of the auditory scene, involving ambient sounds, noises and vocal signals and triggering enhanced brain activations and coupled connectivity in the primary auditory cortex as well as in more secondary areas such as the bilateral TC_lat_ (Fig.7**a**); ii) to focus more specifically on voice signals and therefore clarify communication, the TC_lat_ regions would still be active together with a potentially parallel, anti-coupled network including the posterior TC_lat_ as well as the TC_med_ that would reduce the impact of ambient noise (Fig.7**b**); iii) finally, ‘noise-free’ or ‘noise-reduced’ voice processing would allow for enhanced activity and coupled connectivity between the TC_lat_—especially the mid and anterior subparts—and the IFC (Fig.7**c**), especially the *pars triangularis* to allow for a cognitive evaluation of the voice as well as a decision to be made about this vocal signal—depending on context and task demands. The first step of this model is supported by many previous studies by us^8,11,12,14,16,46^ and others^3,4,6,13,21,27,32,35,36^, and the same is true for the third step of this model in which the IFC is recruited for decisional processes of auditory vocal signals^14,15,46-48^. The second step however still represents a grey area, and the present work aimed at clarifying the role of TC_lat_ and TC_med—_and especially of the bilateral posterior parahippocampal gyrus—in favoring voice processing in noise. These medial regions are indeed at the crossroads of vocal and non-vocal sound processing, with evidence for their distinct involvement in noise^17,19,26,40,41^ as well as in voice or music processing, for instance when emotional content is included^21^. So far, existing literature of voice or speech perception in noise revealed the role of the Medial Olivocochlear Efferent System^2^, brainstem^43,45^, and primary and secondary auditory cortices, which were also reported as core brain regions for voice-in-noise neural development from childhood to adulthood^18^. A recent study pointed to the role of a limbic-cortical network as a basis for separating emotional voices from noise in an auditory scene, interestingly involving subparts of the medial temporal cortex, namely the amygdala^33^. While emotional voices might represent a special case of voice stimuli, these results are still in line with what we observed here and also add more weight to the second step of our model of voice processing in noise. The amygdala and the parahippocampal gyrus—two regions that are next to each other anatomically—could therefore share a role in helping voice perception in noise for emotional and non-emotional content, respectively. This assumption could reinforce the TC_med_-TC_lat_ network hypothesis for noise reduction, but it should be tested scientifically in a dedicated study by using a mix of emotional and non-emotional voice stimuli. Noteworthy is the fact that no study to date highlighted the specific role of the parahippocampal gyrus in voice perception in noise although its involvement in environmental noise processing was previously reported^49^. Interestingly, the parahippocampal gyrus was also repeatedly reported in tinnitus^50-53^, a condition in which patients hear ringing, roaring or buzzing sounds, that are allegedly extremely close to what we would categorize as ‘noise’. The parahippocampal gyrus might therefore play a crucial— and causal—role in the perception, processing and reduction of auditory noise.

At least one open question remains concerning our model and the way it illustrates the results, especially about the mentioned brain networks in step 2 and 3: are these brain networks sequential or parallel? In fact, one could assume that these processes are dealt with in parallel by the brain, since it seems to be a way of functioning especially true for Humans compared to other animals^54^. Our results cannot give a clear answer to this point and while no relevant extrapolation can be made here, we can only say that our results do not exclude either sequential or parallel processing and neural functioning. Future studies should therefore focus on the brain dynamics of voice perception in noise to address this aspect as reported for emotional prosody processing in the past^34^, by the use of electroencephalography or magnetoencephalography for instance. The time-course of this process may also help characterize specific biological markers of the impact of noise exposition on health^23-25^, and this aspect goes beyond communication and is of course of the highest importance for health care.

Several methodological and conceptual limitations should be considered to put our results into perspective and point out what remains to be investigated in the future. First, sample size of Study 3 could have been improved—even though our simulation power calculations gave us optimism concerning effect size in our regions of interest compared to a sample of N=35 or even N=50. Second, ecological validity was not optimal since laboratory studies do not represent real-world experience, especially for concrete topics such as voice-in-noise perception. Other brain networks or regions could therefore be responsible for processing voice signals in noise but their implication could have been attenuated or even annihilated in the laboratory. Third, a direct characterization of the specific acoustical parameters potentially impacting voice perception in noise should be performed, especially using the Mel-Frequency Cepstral Coefficients (MFCC) as a measure of acoustic stimulus organization (‘acoustic entropy’). Fourth, even though we designed Study 3 with task-triggered sparse-sampling fMRI to allow for silent periods during stimulus presentation, we cannot exclude that scanner noise still impacted perception in a differed manner. Fifth, the low temporal resolution of fMRI—in our case 1 second— prevents any reliable testing of sequential or parallel functioning of the observed brain networks. Finally, 3T fMRI does not allow for any assumption or observation of layer-level mechanisms potentially adding up to our results of voice processing in noisy situations. This is especially true for the auditory cortex and/or the parahippocampal gyrus. Future studies should therefore investigate layer-level wholebrain or connectivity data in these regions of interest using for instance high field fMRI at 7T. By doing so, the resulting granularity of research data in this topic would be improved, yielding to more representative brain mechanisms of realistic voice perception.

In conclusion, vocal communication requires humans to filter out—at least partially—sources of noise that would otherwise alter perception and potentially negatively impact social interactions and ultimately communication. We directly manipulated background noise in auditory scenes in which voice signals were presented. High compared to low noise situations altered voice perception and causally relied on a network of regions including the lateral and medial temporal cortex (TC_lat_, TC_med_). Based on our data, we propose a three-step model of voice perception in noise—(1) auditory scene processing, (2) noise reduction, (3) final voice processing and decision. Further research should address the potential parallel functioning of these brain networks and assess whether other regions may be involved in order to draw a clearer picture of functional brain organization related to everyday life communication, especially in the case of voice processing in noisy situations.

## Methods

### Study 1

#### Participants

Ninety-eight right-handed, healthy, either native or highly proficient French-speaking participants (45 male, 53 female, mean age 24.82 years, SD 5.45) were included in this functional magnetic resonance study. All participants were naive to the experimental design and study, had normal or corrected-to-normal vision, normal hearing and no history of psychiatric or neurologic incidents. Participants gave written informed consent for their participation in accordance with ethical and data security guidelines of the University of Geneva. The study was approved by the Ethical Committee of the University of Geneva and was conducted according to the Declaration of Helsinki.

For this study, we did not determine sample size *a priori* since we had a rather large sample already at hand. In the princeps study by Belin and colleagues, reliable and replicated^11,14,16,27^ results were observed with only fourteen participants. We have here in Study 1 about seven times more participants, and this sample size and results are therefore allowing for a very low type I error.

#### Stimuli

Auditory stimuli consisted of sounds from a variety of sources. Vocal stimuli were obtained from 47 speakers: 7 babies, 12 adults, 23 children and 5 older adults. Stimuli included 20 blocks of vocal sounds and 20 blocks of non-vocal sounds. Vocal stimuli within a block could be either speech (33%: words, non-words, foreign language) or non-speech (67%: laughs, sighs, various onomatopoeia). Non-vocal stimuli consisted of natural sounds (14%: wind, streams), animals (29%: cries, gallops), the human environment (37%: cars, telephones, airplanes) or musical instruments (20%: bells, harp, instrumental orchestra). The paradigm, design and stimuli were obtained through the Voice Neurocognition Laboratory website (http://vnl.psy.gla.ac.uk/resources.php). Stimuli were presented through headphones (model S14, Sensimetrics Corporation, Gloucester, MA, USA) at an intensity that was kept constant throughout the experiment (70 dB sound-pressure level). Participants were instructed to actively listen to the sounds. The silent interblock interval was 8 s long.

#### Image acquisition

Structural and functional brain imaging data were acquired by using a 3T scanner (Siemens Trio, Erlangen, Germany) with a 32-channel coil. A magnetization-prepared rapid acquisition gradient echo sequence was used to acquire high-resolution (1 × 1 × 1 mm3) T1-weighted structural images (TR = 1,900 ms, TE = 2.27 ms, TI = 900 ms). Functional images were acquired by using a multislice echo planar imaging sequence (36 transversal slices in descending order, slice thickness 3.2 mm, TR = 2,100 ms, TE = 30 ms, field of view = 205 × 205 mm2, 64 × 64 matrix, flip angle = 90°, bandwidth 1562 Hz/Px).

#### Image analysis

##### Whole-brain analyses

Functional images were analyzed with Statistical Parametric Mapping software (SPM12, Wellcome Trust Centre for Neuroimaging, London, UK). Preprocessing steps included realignment to the first volume of the time series, slice timing, normalization into the Montreal Neurological Institute^55^ (MNI) space using the DARTEL toolbox^56^ and spatial smoothing with an isotropic Gaussian filter of 8 mm full width at half maximum. To remove low-frequency components, we used a high-pass filter with a cutoff frequency of 128 s. Anatomical locations were defined with a standardized coordinate database (Talairach Client, http://www.talairach.org/client.html) by transforming MNI coordinates to match the Talairach space and transforming it back into MNI for display purposes.

A general linear model was used to compute first-level statistics, in which each block was modeled by using a block function and was convolved with the hemodynamic response function, time-locked to the onset of each block. Separate regressors were created for each condition (vocal and non-vocal; "condition" factor). Finally, six motion parameters were included as regressors of no interest to account for movement in the data. The condition regressors were used to compute simple contrasts for each participant, leading to a main effect of vocal and non-vocal at the first-level of analysis ([1 0] for vocal, [0 1] for non-vocal). These simple contrasts were then taken to a flexible factorial second-level analysis in which there were two factors: the "participants" factor (independence set to "yes", variance set to unequal) and the "condition" factor (independence set to "no", variance set to unequal). All neuroimaging activations were thresholded in SPM12 by using a voxel-wise family-wise error (FWE) correction at p<0.05. Whole-brain, second-level analysis results were used to extract and delineate 26 ROIs from the main contrasts (Voice > Non-voice: 19 ROIs; non-vocal>vocal: 7 ROIs) as separate masks (higher accuracy) rather than as spheres around the peak coordinates.

#### Connectivity analyses

##### Seed-to-voxel and seed-to-seed functional and effective connectivity

Functional and effective connectivity analyses were computed by using CONN^30^ (version 22.a) implemented in Matlab 9.0 (The MathWorks, Inc., Natick, MA, USA). Functional and anatomical data were first imported with the SPM import option (automatically loading data from the ‘SPM.mat’ file created in the whole-brain first-level analyses). Following this step, the 26 ROI were entered in the model and seed-to-voxel and seed-to-seed analyses were enabled by using parametric statistics. The following step was denoising, during which data are by default band-pass filtered and detrended and finally artifact corrected (removal of white matter, cerebrospinal fluid, motion, outlier regressors). The last analysis steps are first-level (with SPM regressors, i.e., movement parameters, entered as covariates of no interest) and second-level statistical analyses during which single-subject and group-level analyses are computed, respectively. For each participant and for each condition (vocal; non-vocal), the correlational measures were bivariate and one coefficient was calculated for each seed and target ROI (seed-to-seed analyses) and for each seed and voxels (seed-to-voxel analyses). All analysis steps described were computed by using an explicit mask of the brain to remove any non-brain tissue and the analysis space was kept from the input files on SPM.

Seed-to-voxel analyses were computed with the aim of understanding functional connectivity originating from our ROIs and expanding by increased/reduced functional connectivity with voxels in the whole brain for our contrast of interest (Voice > Non-voice). For seed-to-voxel analysis, all activations are reported at a voxel-wise false-discovery rate (FDR) corrected threshold of p<.01, two-tailed.

On the other hand, seed-to-seed analyses were computed in order to characterize the direct links (increased/reduced functional connectivity) between our extracted ROIs for the contrast of interest (Voice > Non-voice). Results for seed-to-seed analyses are reported at an analysis-level FDR correction for multiple comparisons with a threshold of p<.01, two-tailed.

Finally, seed-to-voxel analyses were computed by using atlas-based labeling implemented in CONN to accurately localize certain subregions of our ROIs, notably to identify which part(s) of the parahippocampal gyri were involved in a negatively connected network, as found in both seed-to-voxel and seed-to-seed analyses mentioned earlier for the Voice > Non-voice contrast. This analysis was computed for several regions of the MTL, notably the amygdalae, hippocampi and parahippocampal gyri. For these seed-to-voxel atlas-based analyses, results are reported at FWE voxel-level correction of p<.01, two-tailed.

### Study 2 & 3

#### Participants

Eighteen (Study 2: 9 male, 9 female, mean age 21.17 years, SD 4.30) and twenty (Study 3: 11 male, 9 female, mean age 22.90 years, SD 3.64) right-handed, healthy, either native or highly proficient French-speaking participants were included in the analyses. The samples of the two studies were completely independent. All participants were naive to the experimental design and study, had normal or corrected-to-normal vision, normal hearing and no history of psychiatric or neurologic incidents. Participants gave written informed consent for their participation in accordance with ethical and data security guidelines of the University of Geneva. The study was approved by the Ethical Committee of the University of Geneva and was conducted according to the Declaration of Helsinki.

Concerning power of Study 2 & 3, we aimed at 80% of power. Using G*Power^57^ software (version 3.1.9.7), we therefore determined that for a medium effect size of 0.5, an alpha of .05 for a two-tailed test, we needed twenty participants to obtain such power. Two participants had to be excluded from Study 2 due to corrupted logfiles while the whole sample was acquired for Study 3. Therefore, after taking this subtraction into account we finally obtained a post-hoc power of 74% instead of 80% for Study 2.

For Study 3, we additionally used the ‘neuRosim’ R package^58^ to simulate nifti format fMRI data according to the specific design of the task—including: the timing of each trial (event-related), the trial sequence (onsets, durations) for each condition in a pseudo-randomized order (no more than three times the same condition in a row). This procedure allowed us to precisely assess fMRI power with N=20. MRI sequence properties were also specified: repetition timing (1 sec), number of scans per session (N=444), field of view, final smoothing of the data (8mm), the specified ROIs (N=10)—including the bilateral posterior STG, MTG, amygdala and parahippocampal gyrus (PHG) with iteratively varying (±3mm) peak coordinates around a given peak voxel—‘runif’ function. Region-specific effect size was also included—ranging from 0.25 (amygdala) to 0.75 (STG/MTG and PHG)—with alpha=.05, power=95%. Noise mixture of type ‘rician’ was also added to better simulate the data. Simulations were computed using non-parametric data generation using 10’000 iterations for a total of 35 simulated participants, exported in 3D nifti format (‘oro.nifti’ R package^59^) and then preprocessed and analyzed in SPM12 with one regressor per condition—conditions included: Highest noise, High noise, Equal, Low noise, Lowest noise—in addition to six movement parameters (Design matrix column number: five conditions + six movement parameters = 11 columns). The results show very good ROI activations at the highest statistical thresholding usually used in publications (p < .05 Family-Wise Error corrected for multiple comparisons) for N=35, N=30 and N=20 (Fig.8**c**, 8**b** and 8**a**, respectively) when contrasting factor modalities at the lowest hierarchical level of the factorial architecture, such as in the comparison between High and Low noise conditions. Simulation methods are state-of-the-art and therefore make for a strong statistical basis for this last study. Twenty participants were therefore included in Study 3, representing the maximum number we could recruit due to financial limitations at this point while still maintaining strong statistical power, as shown by this conservative simulation procedure.

**Fig. 8:**
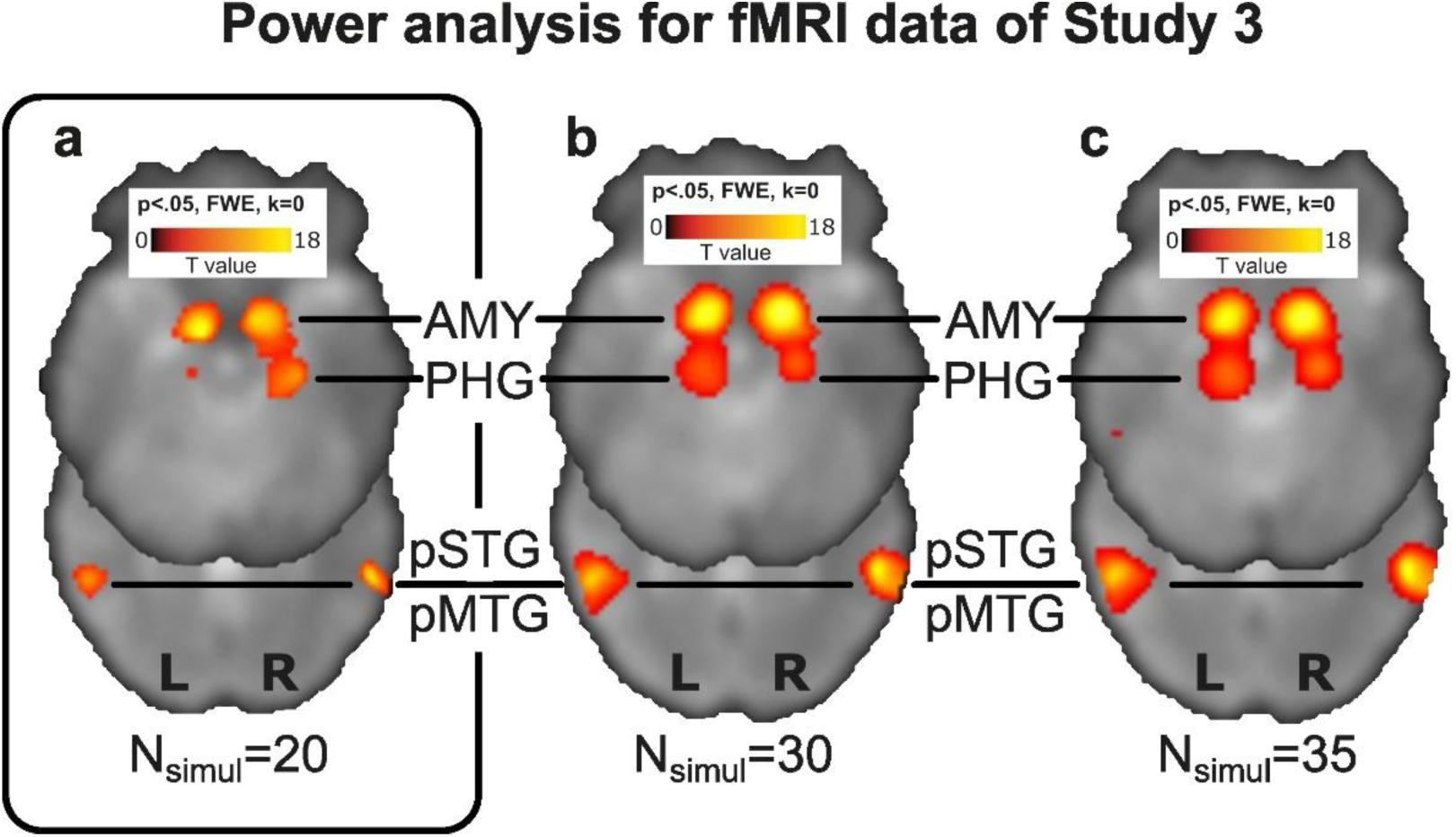
Power analysis using simulated fMRI data, task-specific. The ‘neuRosim’ R package was used to simulate nifti volumes according to the specific parameters, conditions and trial timings of the task of Study 3. Analyses were then performed in SPM12 at the first- and second-level and thresholded at p<.05 Family-Wise Error (FWE) corrected for samples with (**a**) N=20, (**b**) N=30 and (**c**) N=35. Due to financial limitations and because high statistical power was obtained already with N=20, we included twenty participants in Study 3 (**a**, black rounded square). The colorbars represent the t-value, fixed to the highest level, namely the highest t-value obtained in the global maxima of the N=35 sample (T_(34)_=18). N_simul_: simulate sample size; AMY: amygdala; PHG: parahippocampal gyrus; pMTG: posterior division of the middle temporal gyrus; pSTG: posterior division of the superior temporal gyrus; L: left hemisphere; R: right hemisphere.

#### Stimuli

Auditory stimuli consisted of a subset of those used in Study 1, namely stimuli from the Voice Neurocognition Laboratory website (http://vnl.psy.gla.ac.uk/resources.php). Vocal stimuli (47 speakers: 7 babies, 12 adults, 23 children and 5 older adults) were selected to include only those with vocal cords usage, excluding coughs, sighs and throat clearing sounds. Vocal stimuli could therefore be either speech (60%: words, non-words, foreign language) or non-speech (40%: laughs, various onomatopoeia). Non-vocal stimuli consisted of natural sounds (30%: wind, streams) and the human environment (70%: cars, telephones, airplanes). To effectively manipulate the level of background noise we mixed the stimuli by pairs—i.e., each vocal stimulus with each non-vocal stimulus—using Matlab (The Mathworks Inc., Natick, MA, USA) at various ratios of voice-vs-noise intensity (VNI; in dB) creating five conditions (N_trials_=45 each): Highest noise (VNI: 1/16:1), High noise (VNI: 1/8:1), Equal (VNI: 1:1), Low noise (VNI: 1:1/8), Lowest noise (VNI: 1:1/16). Mixed stimuli were saved individually offline and then presented during the task in pseudo-random order (see *Procedure* below) at an intensity that was kept constant throughout the experiment (70 dB sound-pressure level; no statistical difference concerning sound energy between the five conditions, all Ps>.5, Fig.3**b**). Average stimulus duration was 1.5s (±0.5s). Five different lists of mixed stimuli were created, allowing for four rotations within our sample of participants for both studies. Every aspect related to the stimuli that we described in this section is identical for Study 2 & 3. Details about the stimuli are illustrated in Fig.3**b**, including an example of a stimulus across the five different conditions including waveform, spectrogram, intensity and fundamental frequency plots.

#### Procedure

The procedure for Study 2 & 3 was identical (See Fig.3). Participants were presented with a first instruction screen, synthesizing the experimenter’s instructions given beforehand, namely they were instructed to “Listen carefully to each sound and evaluate how easy or difficult it is to clearly perceive the voice (a voice is present in each trial)”. Each trial included a first light grey screen with a black fixation cross during which the stimulus was presented through headphones (Study 2: Sennheiser HD-25 II, Sennheiser electronic SE & Co. KG, Germany; Study 3: model S14, Sensimetrics Corporation, Gloucester, MA, USA) for an average duration of 1.5s (±0.5s), followed by a second light grey screen (duration of 2s) with the fixation cross turning white to indicate to the participant that she/he had to respond using either the ‘1’ (easy to perceive the voice signal) or ‘2’ (difficult to perceive the voice signal) button on the response box (fORP 8 button bimanual curved, Cortech Solutions, Inc., Wilmington, NC, USA; button mapping counter-balanced between participants). A blank screen of average duration 500msec (±200msec) ended the trial sequence. During stimulus presentation, the MRI scanner was silent to allow for clear perception of the sounds, while brain volumes were acquired during the response screen (see the section on MRI data analysis below for more details about this aspect). The task included a total of 225 trials for a total task duration of about 15 minutes. Details about the procedure are illustrated in Fig.3**a**.

#### Behavioral data analysis

As mentioned in the Results section, behavioral results of both studies 2 & 3 illustrate the probability of perceiving a voice in a noisy background and were computed using mixed effects, logistic regression analyses for the response (of-interest) and for the associated reaction times (of-no-interest) in R studio^60^.

For the ‘response’ dependent variable, we used a logistic regression using the lme4 ‘glmer’ package^61^ with ‘conditions’ as fixed effect and participant identity (ID), gender and age as random factors, in addition to stimulus identity (ID) and order. The model formula was the following:

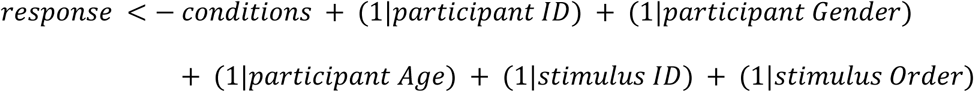

For the ‘reaction times’ dependent variable (RTs), we used a linear regression using the lme4 ‘lmer’ package^1^ with ‘conditions’ as fixed effect and participant identity (ID), gender and age as random factors, in addition to stimulus identity (ID) and order. The model formula was the following:

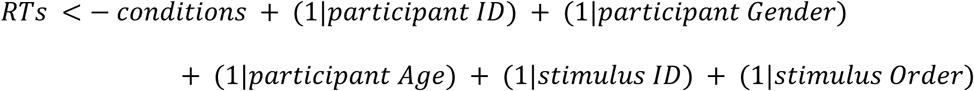

For our planned comparisons, we used a confidence interval of 97.5% and a Bonferroni correction for multiple comparisons. Specific contrasts were computed using the ‘phia’ package^62^. The vector of the main contrast of interest for both dependent variables was the following: [2(Highest noise) 1(High noise) 0(Equal) -1(Low noise) -2(Lowest noise)]. The analyses were identical for Study 2 & 3.

#### Image acquisition

Structural and functional brain imaging data were acquired by using a 3T scanner (Siemens Trio, Erlangen, Germany) with a 32-channel coil. A magnetization-prepared rapid acquisition gradient echo sequence was used to acquire high-resolution (1 × 1 × 1mm3) T1-weighted structural images (TR = 1,900 ms, TE = 2.27 ms, TI = 900 ms). Functional images were acquired using two consecutive 1s volumes post stimulus presentation (during the response screen) in a sparse manner by using a multislice echo planar imaging sequence (56 transversal slices in descending order with 2.5mm isotropic voxels, TR = 1000 ms, TE = 30 ms, field of view = 205 × 205 mm2, 84 × 84 matrix, flip angle = 64°). A total of 444 volumes were acquired for each participant of Study 3.

#### Image analysis

##### Whole-brain analyses

Functional images were analyzed with Statistical Parametric Mapping software (SPM12, Wellcome Trust Centre for Neuroimaging, London, UK). Preprocessing steps included realignment to the first volume of the time series, slice timing, normalization into the Montreal Neurological Institute^55^ (MNI) space and spatial smoothing with an isotropic Gaussian filter of 8mm full width at half maximum. To remove low-frequency components, we used a high-pass filter with a cutoff frequency of 128s. Autocorrelation estimation was set to ‘FAST’ since our MRI repetition time was short (1s).

A general linear model was used to compute first-level statistics, in which each event was modeled by using a convolution with the ‘finite impulse response’ function (fixed duration of 2s), time-locked to the onset of each event. Separate regressors were created for each condition (Highest noise, High noise, Equal, Low noise, Lowest noise; "condition" factor). Finally, six motion parameters were included as regressors of no interest to account for movement in the data. The condition regressors were used to compute simple contrasts for each participant, leading to a main effect of each condition. These simple contrasts were then taken to a flexible factorial second-level analysis in which there were two factors: the "participants" factor (independence set to "yes", variance set to unequal) and the "condition" factor (independence set to "no", variance set to unequal). All neuroimaging activations were thresholded in SPM12 by using a voxel-wise correction for multiple comparisons of p<.05 FDR.

##### Functional and effective connectivity analyses

Functional and effective connectivity analyses (except for dynamic causal modelling, see specific section below) were computed by using the CONN toolbox^30^ (version 22.a) implemented in Matlab 9.0 (The MathWorks, Inc., Natick, MA, USA). Analyses included: seed-to-voxel functional and effective connectivity, functional connectivity multivariate pattern analysis (fc-MVPA).

Functional and anatomical data were first imported with the SPM import option. Following this step, seed-to-voxel analyses were enabled by using parametric statistics. The following step was denoising, during which data are by default band-pass filtered and detrended and finally artifact corrected (removal of white matter, cerebrospinal fluid, motion, outlier regressors). The last analysis steps are first-level (with SPM regressors, i.e., movement parameters, entered as covariates of no interest) and second-level statistical analyses during which single-subject and group-level analyses are computed, respectively.

##### Seed-to-voxel and seed-to-seed functional and effective connectivity analyses

For each participant and for each condition, measures were bivariate (correlation for functional connectivity, regression for effective connectivity) and one coefficient was calculated for each seed (seed-to-seed analyses) and each voxel (seed-to-voxel analyses). All analysis steps described were computed by using an explicit mask of the brain to remove any non-brain tissue and the analysis space was kept from the input files on SPM.

These analyses were computed using generalized psychophysiological interaction (gPPI) maps. These maps represent the level of task-modulated ‘effective’ connectivity between a seed and another seed (seed-to-seed) or every voxels in the brain (seed-to-voxel)—namely, changes in functional association strength covarying with an experimental factor. gPPI is computed using a separate multiple regression model for each target seed/voxel timeseries. For these analyses, seeds were labelled by using the atlas implemented in the CONN toolbox, with the addition of bilateral pMTG and PHG ROIs extracted from Study 1 (see Fig.1**d**-**f**) as binary masks. We chose to include these ROIs to investigate the level of similarity the connectivity patterns of processing noise alone (Study 1) as opposed to processing voice with background noise (Study 3). For these analyses, all activations are reported at a voxel-wise corrected threshold of p<.05 FDR, two-tailed.

##### Voxel-level network analyses: fc-MVPA

Multivariate correlation maps represent for each voxel the number of most salient spatial features of the seed-based connectivity maps seeded at this very same voxel. They are defined by using a Singular Value Decomposition^63,64^ separately for each seed voxel, of the patterns of seed-based correlations across all participants. The motivation behind the use of this additional connectivity measure was to extend and *a minima* compare univariate to multivariate connectivity results. A homogeneity between these two distinct types of analyses gives even more strength to the data and provides valuable coherence. All activations are reported at a voxel-wise corrected threshold of p<.05 FDR, two-tailed.

##### Dynamic causal modelling (DCM)

DCM—a directed, effective connectivity measure—was performed by following the steps recommended by Friston^28,29^. Volumes of interest were first extracted in SPM12 as actual activated clusters—not spheres or squares, including: the right posterior middle temporal gyrus obtained by running the fc-MVPA analysis (Fig.5**a**) mentioned above (peak MNI *xyz* 62,-16,-12); the right parahippocampal gyrus activity resulting from contrasting non-vocal to vocal stimuli (Fig.1**e**, Study 1, peak MNI *xyz* 26,-30,-18) overlapping with the same region in Study 3 (Fig.4**c**); the left posterior parahippocampal gyrus observed in Study 3 gPPIs (Fig.5**b**) when contrasting high noise to low noise vocal discrimination (peak MNI *xyz* -22,-26,-12). First level analyses were then computed per participant by creating three families of models to be tested for the two High noise conditions: the ‘MTG’ models family (Family 1), in which connections originated from the right pMTG to the parahippocampus (Fig.S8**b**); the ‘PHG inputs’ models family (Family 2), in which the connections don’t originate from the pMTG but enter directly the PHGs or their inter-connectivity (FigS8**c**); the last family includes ‘MTG and PHG input’ models (Family 3), in which the connections enter either between the MTG and PHGs or both the MTG and PHGs and their connections (Fig.S8**d**). These families and models were computed in relation with relevant literature^4,18,19,33,41,44,45^ and our task results (Study 3, Fig.4 and Fig.5). Next, fixed effects (FFX), second-level Bayesian model selection (BMS) analyses were computed for our sample of participants. We chose FFX because we were interested in patterns of directed effective connectivity common to our participants, not to differences between them. These analyses revealed Family 3 as the winning family with model 8 (inputs to the right pMTG, both PHGs and to the directed connections from the pMTG to the PHGs) as the overall winning model (posterior probability >.99, Fig.6 and Fig.S8**d**), see the results section.

## Acknowledgements

The authors acknowledge the Swiss National Science Foundation (SNSF project no 105314_124572/1) and the Swiss National Competence Center in Research in affective sciences (51NF40-104897 – DG). The authors also thank Bruno Bonnet, Valérie Milesi-Sterck and Julie Péron, who helped acquire the neuroimaging data as well as Pascal Belin for his insights into this work. LC designed and created the task of Study 2 & 3, programmed all tasks, collected the behavioral and neuroimaging data, analyzed the data, created and edited the figures and wrote and edited the manuscript. ES and DVD helped in the specific design and computation of the connectivity analyses. SF acquired part of the data and helped design the dynamic causal modeling analyses. DG designed the studies, especially Study 2 & 3 and funded all studies. All authors reviewed and edited the manuscript.

## Author statement

The authors declare no competing interests whatsoever.

## Data statement

The data and codes will be made available on the open YARETA repository once the article is published.

## Notes

### Competing Interest Statement

The authors have declared no competing interest.

## References

1 Nitecki, D. V. & Nitecki, M. H. Origins of anatomically modern humans. (Springer Science & Business Media, 2013).

2 Arbib, M. A. From monkey-like action recognition to human language: An evolutionary framework for neurolinguistics. Behavioral and brain sciences 28, 105–124 (2005).

3 Belin, P., Zatorre, R. J., Lafaille, P., Ahad, P. & Pike, B. Voice-selective areas in human auditory cortex. Nature 403, 309–312 (2000).

4 Pernet, C. R. et al. The human voice areas: Spatial organization and inter-individual variability in temporal and extra-temporal cortices. Neuroimage 119, 164–174 (2015).

5 Ethofer, T. et al. Functional responses and structural connections of cortical areas for processing faces and voices in the superior temporal sulcus. Neuroimage 76, 45–56 (2013).

6 Fecteau, S., Armony, J. L., Joanette, Y. & Belin, P. Is voice processing species-specific in human auditory cortex? An fMRI study. Neuroimage 23, 840–848 (2004).

7 Petkov, C. I. et al. A voice region in the monkey brain. Nature neuroscience 11, 367–374 (2008).

8 Ceravolo, L., Frühholz, S., Pierce, J., Grandjean, D. & Péron, J. Basal ganglia and cerebellum contributions to vocal emotion processing as revealed by high-resolution fMRI. Scientific reports 11, 1–15 (2021).

9 Frühholz, S. & Ceravolo, L. The neural network underlying the processing of affective vocalizations. The Oxford handbook of voice perception, 431 (2018).

10 Kreifelts, B., Ethofer, T., Grodd, W., Erb, M. & Wildgruber, D. Audiovisual integration of emotional signals in voice and face: an event-related fMRI study. Neuroimage 37, 1445–1456 (2007).

11 Ceravolo, L., Frühholz, S. & Grandjean, D. Proximal vocal threat recruits the right voice-sensitive auditory cortex. Social cognitive and affective neuroscience 11, 793–802 (2016).

12 Ceravolo, L., Frühholz, S. & Grandjean, D. Modulation of auditory spatial attention by angry prosody: an fMRI auditory dot-probe study. Frontiers in neuroscience 10, 216 (2016).

13 Ethofer, T., Van De Ville, D., Scherer, K. & Vuilleumier, P. Decoding of emotional information in voice-sensitive cortices. Current Biology 19, 1028–1033 (2009).

14 Frühholz, S., Ceravolo, L. & Grandjean, D. Specific brain networks during explicit and implicit decoding of emotional prosody. *Cerebral Cortex*, bhr184 (2011).

15 Frühholz, S. & Grandjean, D. Towards a fronto-temporal neural network for the decoding of angry vocal expressions. Neuroimage 62, 1658–1666 (2012).

16 Grandjean, D. et al. The voices of wrath: brain responses to angry prosody in meaningless speech. Nature neuroscience 8, 145–146 (2005).

17 Schnider, A., Benson, D. F., Alexander, D. N. & Schnider-Klaus, A. Non-verbal environmental sound recognition after unilateral hemispheric stroke. Brain 117, 281–287 (1994).

18 Vander Ghinst, M., et al. Cortical tracking of speech-in-noise develops from childhood to adulthood. Journal of Neuroscience 39, 2938–2950 (2019).

19 Lewis, J. W. et al. Human brain regions involved in recognizing environmental sounds. Cerebral cortex 14, 1008–1021 (2004).

20 Frühholz, S. & Grandjean, D. Amygdala subregions differentially respond and rapidly adapt to threatening voices. Cortex 49, 1394–1403 (2013).

21 Frühholz, S., Trost, W. & Grandjean, D. The role of the medial temporal limbic system in processing emotions in voice and music. Progress in neurobiology 123, 1–17 (2014).

22 Von Kriegstein, K., Kleinschmidt, A., Sterzer, P. & Giraud, A.-L. Interaction of face and voice areas during speaker recognition. Journal of Cognitive Neuroscience 17, 367–376 (2005).

23 Daiber, A. et al. Environmental noise induces the release of stress hormones and inflammatory signaling molecules leading to oxidative stress and vascular dysfunction—Signatures of the internal exposome. Biofactors 45, 495–506 (2019).

24 Münzel, T., Sørensen, M. & Daiber, A. Transportation noise pollution and cardiovascular disease. Nature Reviews Cardiology 18, 619–636 (2021).

25 Jafari, Z., Kolb, B. E. & Mohajerani, M. H. Chronic traffic noise stress accelerates brain impairment and cognitive decline in mice. Experimental neurology 308, 1–12 (2018).

26 Arjunan, A. & Rajan, R. Noise and brain. Physiology & Behavior 227, 113136 (2020).

27 Belin, P., Fecteau, S. & Bedard, C. Thinking the voice: neural correlates of voice perception. Trends in cognitive sciences 8, 129–135 (2004).

28 Friston, K. et al. Psychophysiological and modulatory interactions in neuroimaging. Neuroimage 6, 218–229 (1997).

29 Friston, K. J., Harrison, L. & Penny, W. Dynamic causal modelling. Neuroimage 19, 1273–1302 (2003).

30 Whitfield-Gabrieli, S. & Nieto-Castanon, A. Conn: a functional connectivity toolbox for correlated and anticorrelated brain networks. Brain connectivity 2, 125–141 (2012).

31 Ethofer, T. et al. Emotional voice areas: anatomic location, functional properties, and structural connections revealed by combined fMRI/DTI. *Cerebral Cortex*, bhr113 (2011).

32 Kriegstein, K. V. & Giraud, A.-L. Distinct functional substrates along the right superior temporal sulcus for the processing of voices. Neuroimage 22, 948–955 (2004).

33 Swanborough, H., Staib, M. & Frühholz, S. Neurocognitive dynamics of near-threshold voice signal detection and affective voice evaluation. Science advances 6, eabb3884 (2020).

34 Schirmer, A. & Kotz, S. A. Beyond the right hemisphere: brain mechanisms mediating vocal emotional processing. Trends in cognitive sciences 10, 24–30 (2006).

35 Wildgruber, D., Pihan, H., Ackermann, H., Erb, M. & Grodd, W. Dynamic brain activation during processing of emotional intonation: influence of acoustic parameters, emotional valence, and sex. Neuroimage 15, 856–869 (2002).

36 Latinus, M. & Belin, P. Human voice perception. Current Biology 21, R143–R145 (2011).

37 Iseli, M., Shue, Y.-L. & Alwan, A. Age, sex, and vowel dependencies of acoustic measures related to the voice sourcea). The Journal of the Acoustical Society of America 121, 2283–2295 (2007).

38 Hall, J. A., Gunnery, S. D., Letzring, T., Carney, D. R. & Colvin, C. R. Accuracy of Judging Affect and Accuracy of Judging Personality: How and When Are They Related? Journal of personality (2016).

39 McAleer, P., Todorov, A. & Belin, P. How do you say ‘Hello’? Personality impressions from brief novel voices. PloS one 9, e90779 (2014).

40 Jääskeläinen, I. P. et al. Human posterior auditory cortex gates novel sounds to consciousness. Proceedings of the National Academy of Sciences of the United States of America 101, 6809–6814 (2004).

41 Tomasino, B. et al. Identifying environmental sounds: a multimodal mapping study. Frontiers in human neuroscience 9 (2015).

42 Mishra, S. K. & Lutman, M. E. Top-down influences of the medial olivocochlear efferent system in speech perception in noise. PloS one 9, e85756 (2014).

43 Parbery-Clark, A., Strait, D. & Kraus, N. Context-dependent encoding in the auditory brainstem subserves enhanced speech-in-noise perception in musicians. Neuropsychologia 49, 3338–3345 (2011).

44 Scott, S. K., Blank, C. C., Rosen, S. & Wise, R. J. Identification of a pathway for intelligible speech in the left temporal lobe. Brain 123, 2400–2406 (2000).

45 Song, J. H., Skoe, E., Banai, K. & Kraus, N. Perception of speech in noise: neural correlates. Journal of cognitive neuroscience 23, 2268–2279 (2011).

46 Grandjean, D. Brain networks of emotional prosody processing. Emotion Review 13, 34–43 (2021).

47 Ceravolo, L., Debracque, C., Pool, E., Gruber, T. & Grandjean, D. Frontal mechanisms underlying primate calls recognition by humans. Cerebral Cortex Communications 4, tgad019 (2023).

48 Dricu, M., Ceravolo, L., Grandjean, D. & Frühholz, S. Biased and unbiased perceptual decision-making on vocal emotions. Scientific reports 7, 16274 (2017).

49 Milne, J. L., Arnott, S. R., Kish, D., Goodale, M. A. & Thaler, L. Parahippocampal cortex is involved in material processing via echoes in blind echolocation experts. Vision research 109, 139–148 (2015).

50 Chen, J. et al. Sensorineural hearing loss affects functional connectivity of the auditory Cortex, parahippocampal gyrus and Inferior prefrontal gyrus in tinnitus patients. Frontiers in Neuroscience 16, 816712 (2022).

51 De Ridder, D., Friston, K., Sedley, W. & Vanneste, S. A parahippocampal-sensory Bayesian vicious circle generates pain or tinnitus: a source-localized EEG study. Brain communications 5, fcad132 (2023).

52 De Ridder, D. & Vanneste, S. Targeting the parahippocampal area by auditory cortex stimulation in tinnitus. Brain stimulation 7, 709–717 (2014).

53 Elgoyhen, A. B., Langguth, B., De Ridder, D. & Vanneste, S. Tinnitus: perspectives from human neuroimaging. Nature Reviews Neuroscience 16, 632–642 (2015).

54 Griffa, A. et al. Evidence for increased parallel information transmission in human brain networks compared to macaques and male mice. Nature Communications 14, 8216 (2023).

55 Collins, D. L., Neelin, P., Peters, T. M. & Evans, A. C. Automatic 3D intersubject registration of MR volumetric data in standardized Talairach space. Journal of computer assisted tomography 18, 192–205 (1994).

56 Ashburner, J. A fast diffeomorphic image registration algorithm. Neuroimage 38, 95–113 (2007).

57 Kang, H. Sample size determination and power analysis using the G* Power software. Journal of educational evaluation for health professions 18 (2021).

58 Welvaert, M., Durnez, J., Moerkerke, B., Verdoolaege, G. & Rosseel, Y. neuRosim: An R package for generating fMRI data. Journal of Statistical Software 44, 1–18 (2011).

59 Whitcher, B., Schmid, V. J. & Thorton, A. Working with the DICOM and NIfTI Data Standards in R. Journal of Statistical Software 44, 1–29 (2011).

60 RStudio, T. (PBC Boston, MA, USA, 2020).

61 Team, R. C. (R Foundation for Statistical Computing. https://www.R-project.org, 2018).

62 De Rosario-Martinez, H., Fox, J., Team, R. C. & De Rosario-Martinez, M. H. Package ‘phia’. CRAN Repos. Retrieved 1, 2015 (2015).

63 Strang, G. The fundamental theorem of linear algebra. The American Mathematical Monthly 100, 848–855 (1993).

64 Wall, M. E., Rechtsteiner, A. & Rocha, L. M. in A practical approach to microarray data analysis 91–109 (Springer, 2003).

