## Supplementary figures for "Functional and causal neural mechanisms of human voice perception in noisy situations"

**Supplementary material**

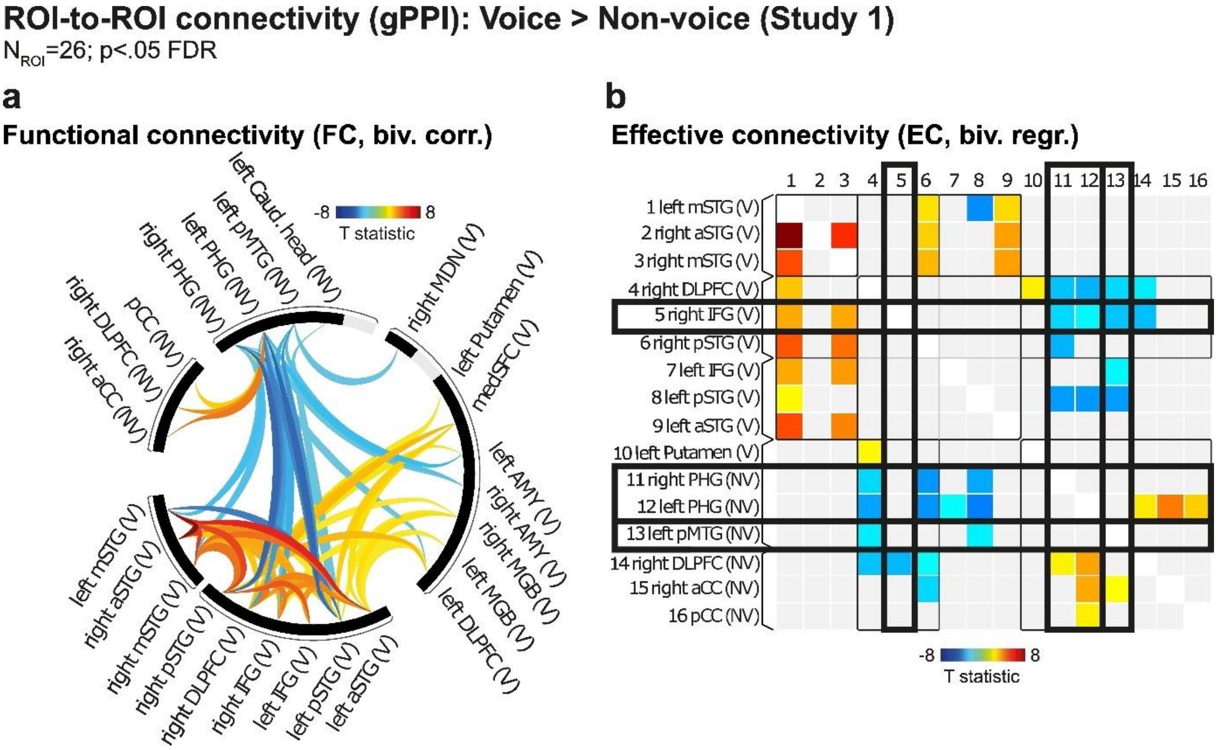


**Fig.S1: Connectivity measures using generalized psychophysiological interaction analyses between ROIs in Study 1 for Vocal > Non-vocal, p<.05 FDR.** (**a**) Functional connectivity using bivariate correlations between our 26 ROIs extracted from Study 1. (**b**) Undirected effective connectivity using bivariate regressions between our 26 ROIs extracted from Study 1. Relations between our ROIs are outlined in black. Colormaps represent T statistics. Thresholding: p<.05 FDR corrected.

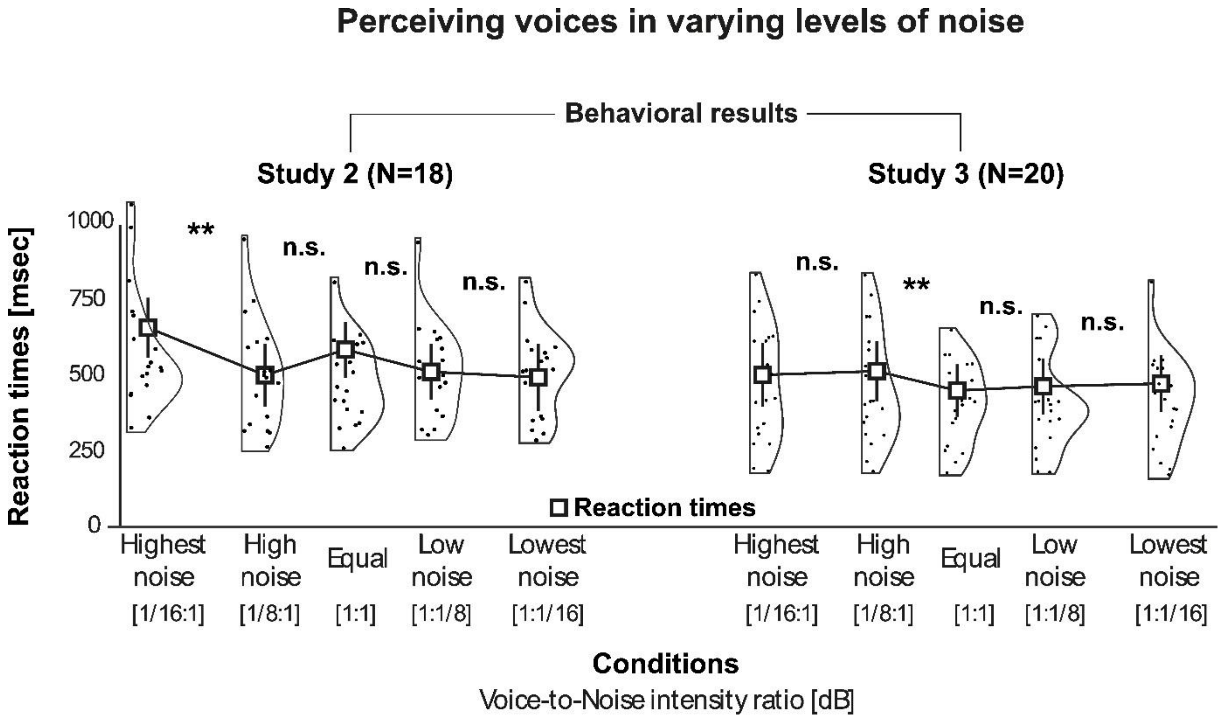


**Fig.S2: Reaction times data when assessing voice perception in noise (Study 2 & 3).** Behavioral results of both studies illustrating response reaction times when perceiving a voice in a noisy background (linear regression analysis, circles indicate individual values, half-violin curves the distribution of the data), Y axis. (left panel: Study 2, N=18; right panel: Study 3, N=20). X axis: conditions with voice-to-noise ratio, in decibel (dB). Error bars represent the standard error of the mean. n.s.: not significant; $^{**}p<.01$.

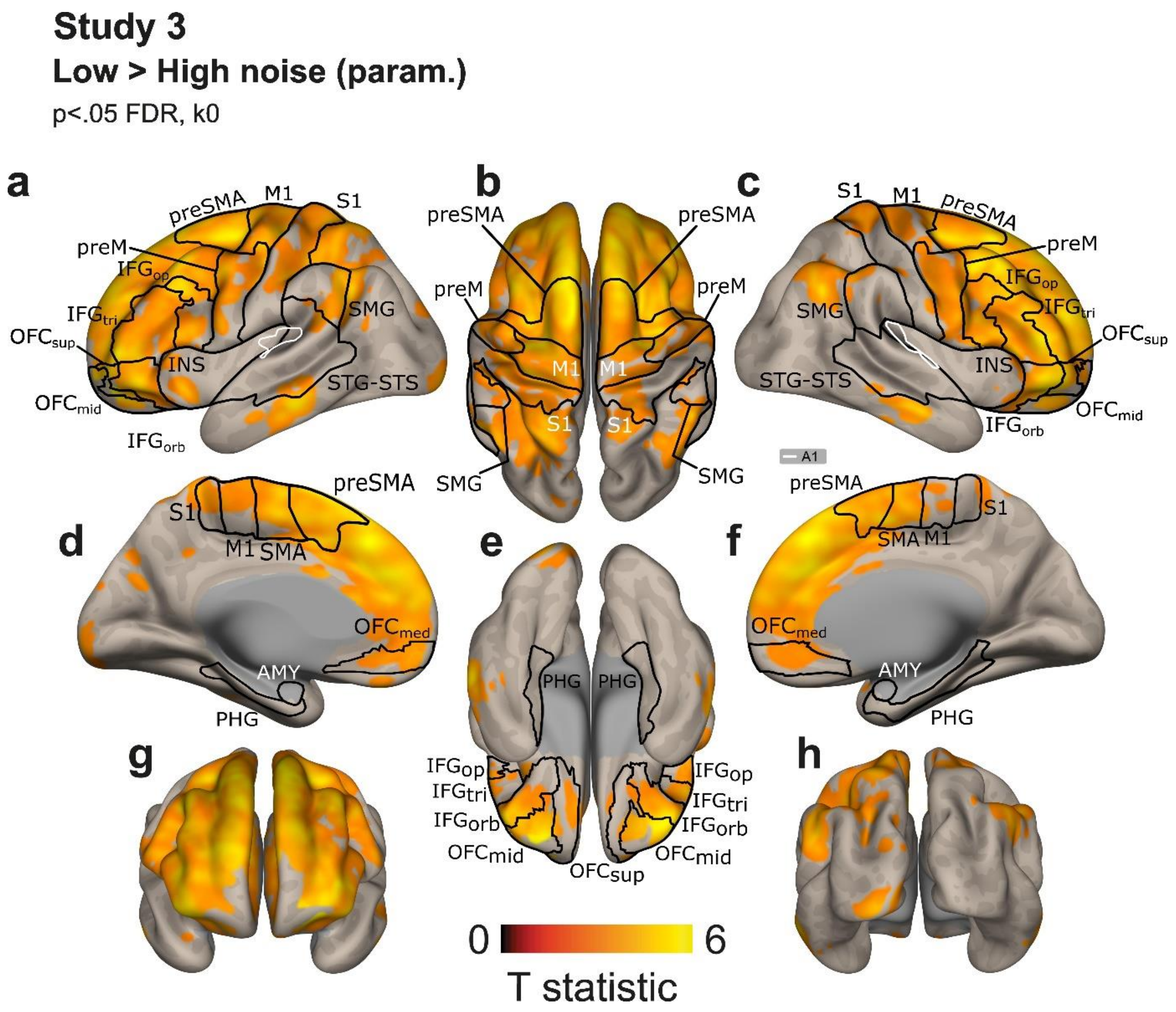


**Fig.S3: Voice perceived in Low > High noise (Study 3).** Whole-brain neuroimaging results highlighting vast frontal activations (**a,b,c,e,g**) as well as medial activity (**d,f**) and posterior activations (**h**). Statistically significant clusters are displayed on a normalized template at a threshold of $p<.05$, corrected for multiple comparisons at the voxel level (False Discovery Rate; FDR). The color bars illustrate the ‘t’ statistical values. A1: primary auditory cortex; IFG: inferior frontal gyrus; INS: insula; MTG: middle temporal gyrus; STG: superior temporal gyrus; STS: superior temporal sulcus; PHG: parahippocampal gyrus; AMY: amygdala; preSMA: pre-supplementary motor area; preM: premotor cortex; M1: primary motor cortex; S1: primary somatosensory cortex; SMG: supramarginal gyrus; OFC: orbitofrontal cortex; TVAs: temporal voice areas. Suffixes: orb, pars orbitalis; tri, pars triangularis; op, pars opercularis; sup, superior; mid, middle; med, medial.

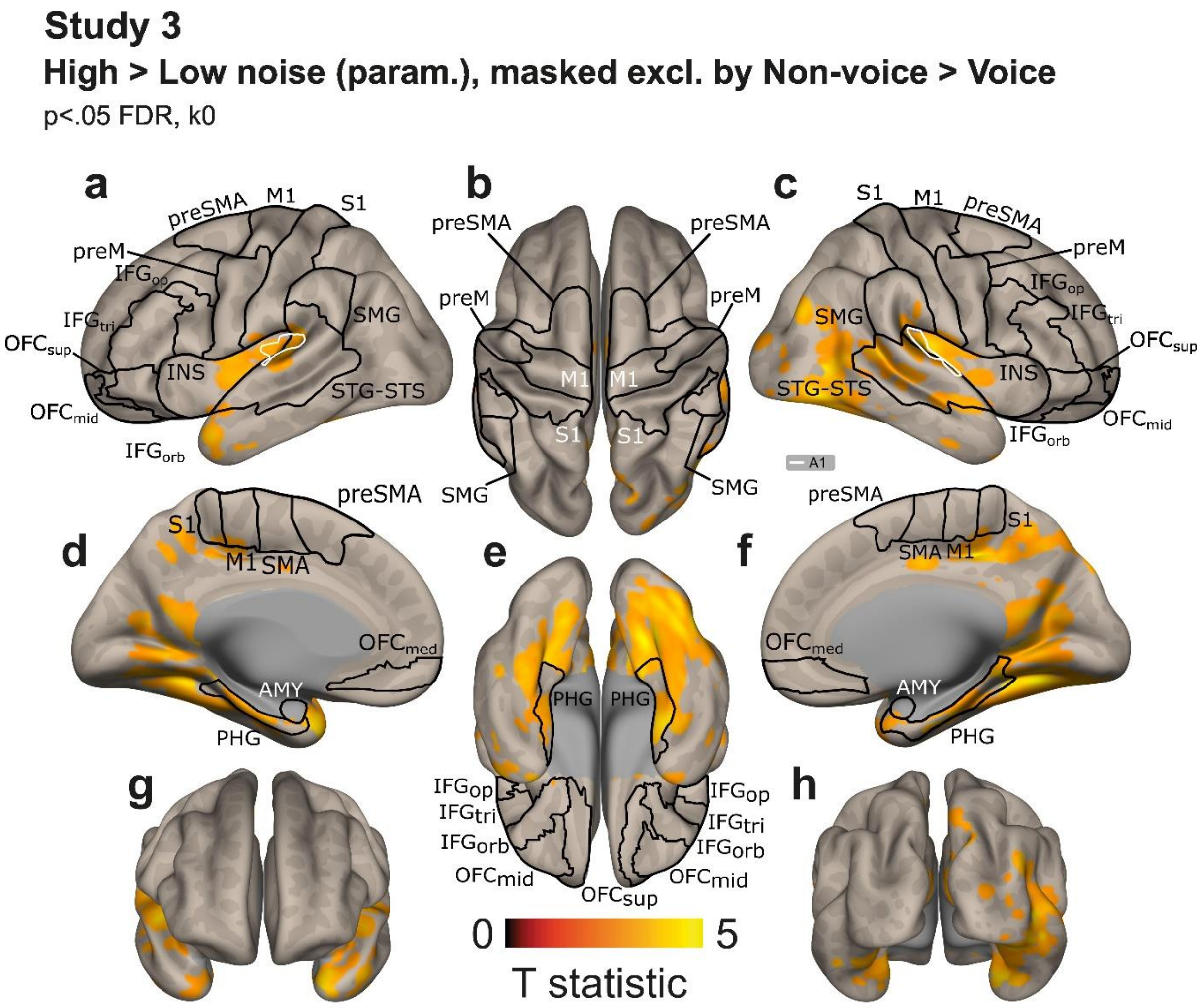


**Fig.S4: Voice perceived in High > Low noise (Study 3) masked exclusive of Non-vocal > Vocal results (Study 2).** Whole-brain neuroimaging results highlighting vast frontal activations (**a,b,c,e,g**) as well as medial activity (**d,f**) and posterior activations (**h**). Statistically significant clusters are displayed on a normalized template at a threshold of $p<.05$, corrected for multiple comparisons at the voxel level (False Discovery Rate; FDR). The color bars illustrate the ‘t’ statistical values. A1: primary auditory cortex; IFG: inferior frontal gyrus; INS: insula; MTG: middle temporal gyrus; STG: superior temporal gyrus; STS: superior temporal sulcus; PHG: parahippocampal gyrus; AMY: amygdala; preSMA: pre-supplementary motor area; preM: premotor cortex; M1: primary motor cortex; S1: primary somatosensory cortex; SMG: supramarginal gyrus; OFC: orbitofrontal cortex; TVAs: temporal voice areas. Suffixes: orb, pars orbitalis; tri, pars triangularis; op, pars opercularis; sup, superior; mid, middle; med, medial.

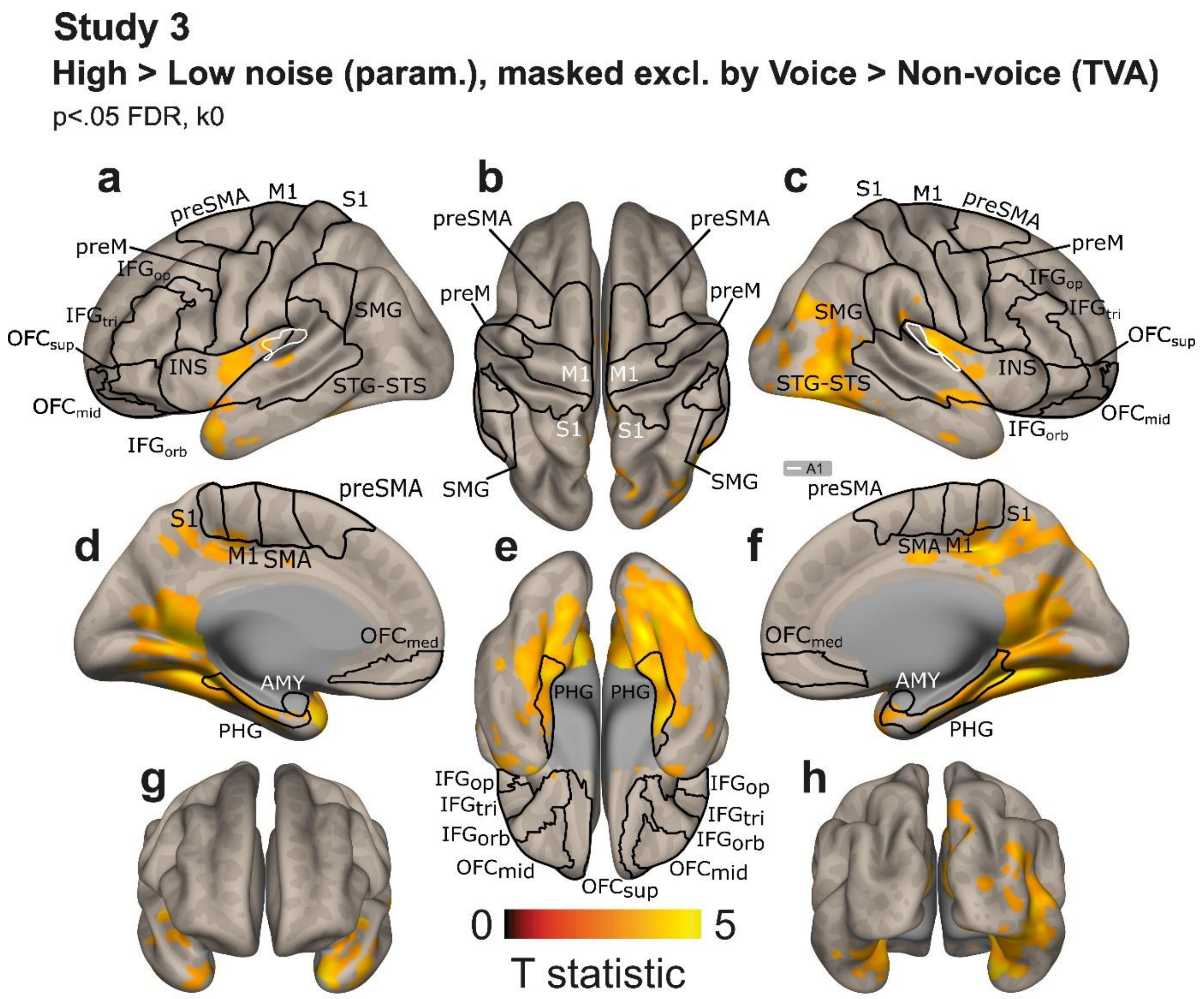


**Fig.S5: Voice perceived in High > Low noise (Study 3) masked exclusive of Vocal > Non-vocal results (Study 2).** Whole-brain neuroimaging results highlighting vast frontal activations (**a,b,c,e,g**) as well as medial activity (**d,f**) and posterior activations (**h**). Statistically significant clusters are displayed on a normalized template at a threshold of $p<.05$, corrected for multiple comparisons at the voxel level (False Discovery Rate; FDR). The color bars illustrate the 't' statistical values. A1: primary auditory cortex; IFG: inferior frontal gyrus; INS: insula; MTG: middle temporal gyrus; STG: superior temporal gyrus; STS: superior temporal sulcus; PHG: parahippocampal gyrus; AMY: amygdala; preSMA: pre-supplementary motor area; preM: premotor cortex; M1: primary motor cortex; S1: primary somatosensory cortex; SMG: supramarginal gyrus; OFC: orbitofrontal cortex; TVAs: temporal voice areas. Suffixes: orb, pars orbitalis; tri, pars triangularis; op, pars opercularis; sup, superior; mid, middle; med, medial.

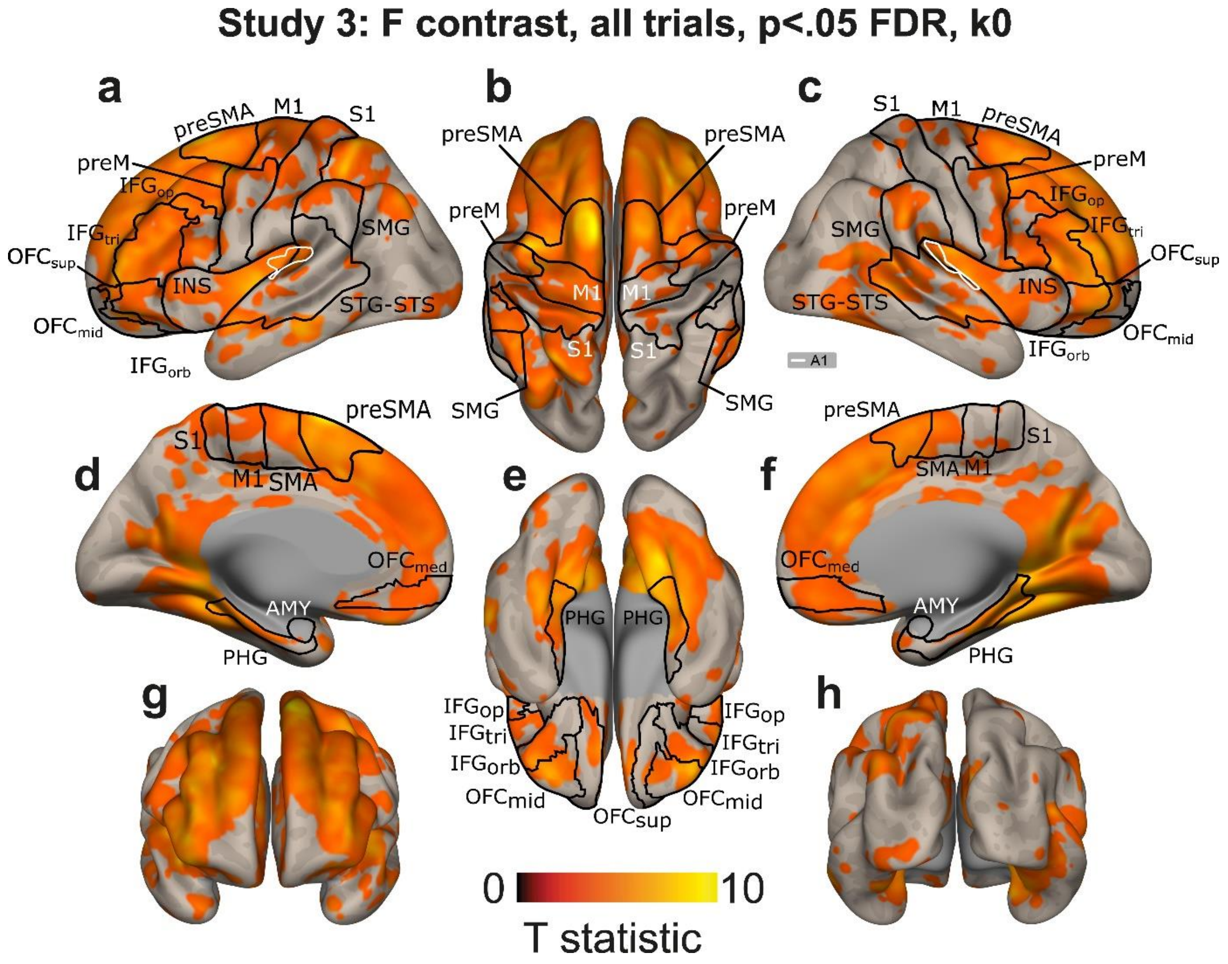


**Fig.S6: Voice in noise (all trials, independent of the participants' response), F-test 'any effect' (Study 3).** Whole-brain neuroimaging results highlighting vast frontal activations (**a,b,c,e,g**) as well as medial activity (**d,f**) and posterior activations (**h**). Statistically significant clusters are displayed on a normalized template at a threshold of $p<.05$, corrected for multiple comparisons at the voxel level (False Discovery Rate; FDR). The color bars illustrate the 't' statistical values. A1: primary auditory cortex; IFG: inferior frontal gyrus; INS: insula; MTG: middle temporal gyrus; STG: superior temporal gyrus; STS: superior temporal sulcus; PHG: parahippocampal gyrus; AMY: amygdala; preSMA: pre-supplementary motor area; preM: premotor cortex; M1: primary motor cortex; S1: primary somatosensory cortex; SMG: supramarginal gyrus; OFC: orbitofrontal cortex; TVAs: temporal voice areas. Suffixes: orb, pars orbitalis; tri, pars triangularis; op, pars opercularis; sup, superior; mid, middle; med, medial.

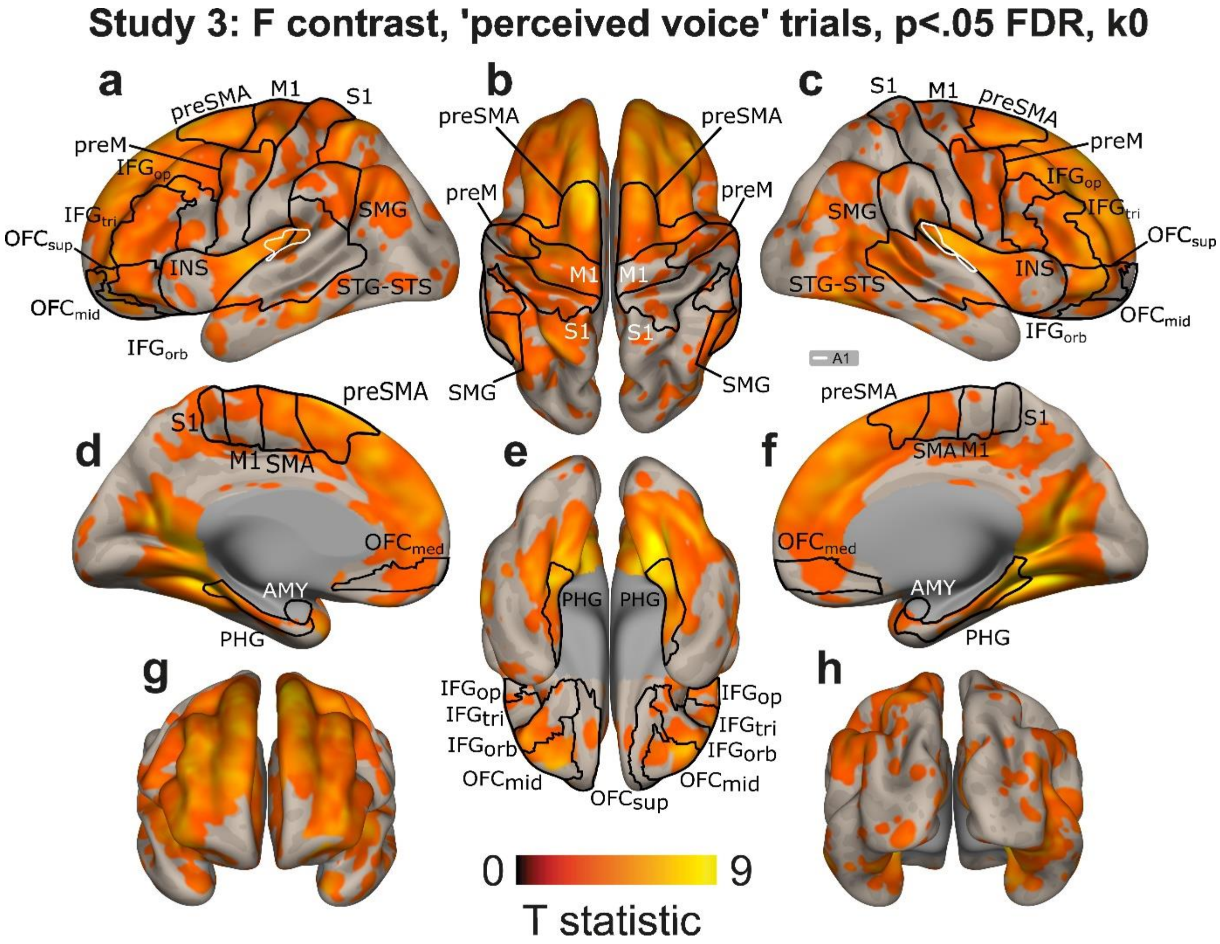


**Fig.S7: Voice perceived in noise (dependent of the participants' response: voices 'easy' to perceive), F-test 'any effect' (Study 3).** Whole-brain neuroimaging results highlighting vast frontal activations (**a,b,c,e,g**) as well as medial activity (**d,f**) and posterior activations (**h**). Statistically significant clusters are displayed on a normalized template at a threshold of $p<.05$, corrected for multiple comparisons at the voxel level (False Discovery Rate; FDR). The color bars illustrate the 't' statistical values. A1: primary auditory cortex; IFG: inferior frontal gyrus; INS: insula; MTG: middle temporal gyrus; STG: superior temporal gyrus; STS: superior temporal sulcus; PHG: parahippocampal gyrus; AMY: amygdala; preSMA: pre-supplementary motor area; preM: premotor cortex; M1: primary motor cortex; S1: primary somatosensory cortex; SMG: supramarginal gyrus; OFC: orbitofrontal cortex; TVAs: temporal voice areas. Suffixes: orb, pars orbitalis; tri, pars triangularis; op, pars opercularis; sup, superior; mid, middle; med, medial.

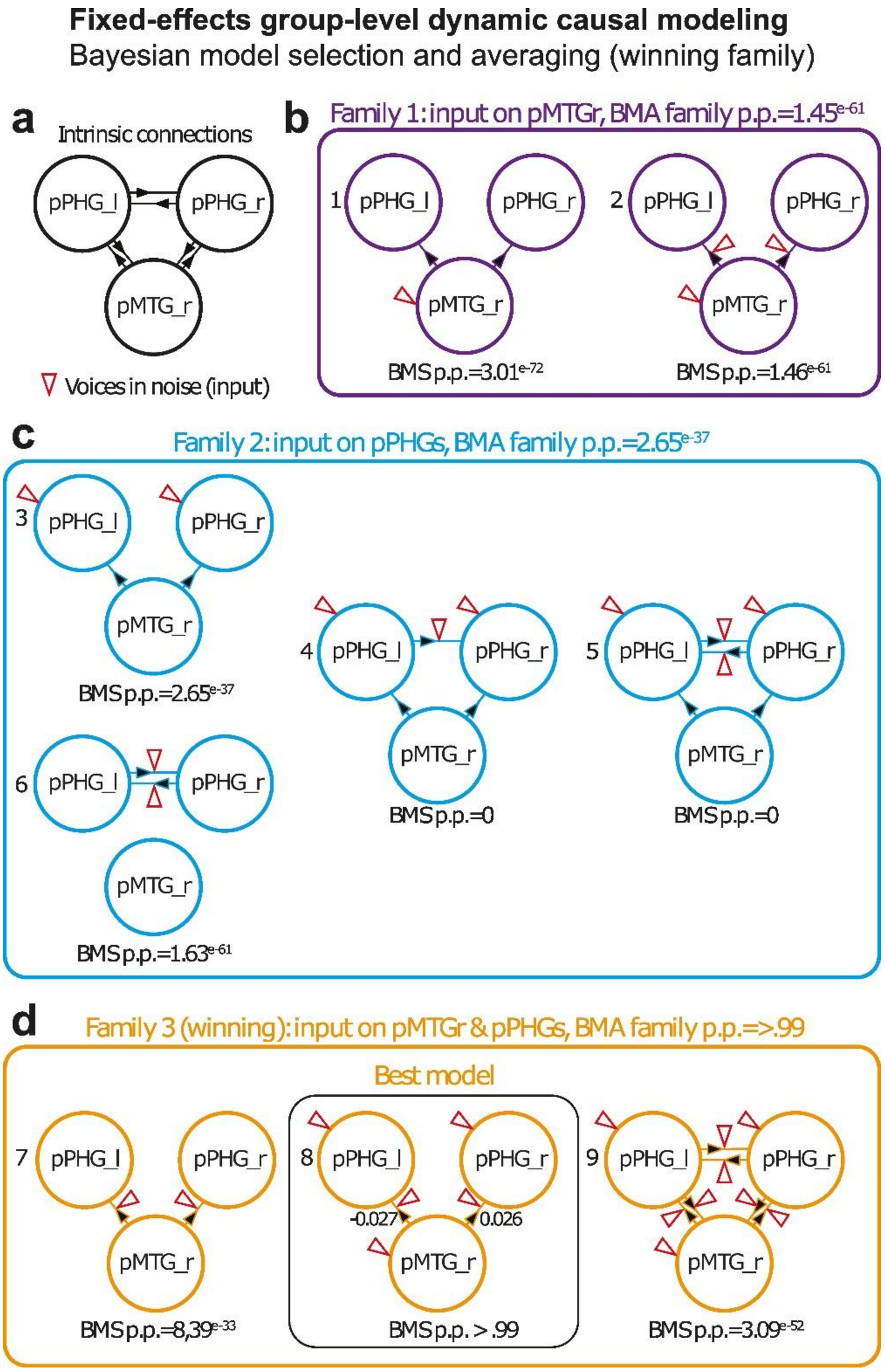


**Fig.S8: Effective, directed fixed-effect group-level connectivity underlying voice processing in noise using dynamic causal modelling (DCM).** DCM analysis was used on nine models and three families, all containing the right posterior middle temporal gyrus (pMTG) and the bilateral posterior parahippocampal gyrus (pPHG). Families and models are detailed in Fig.S8. (**a**) Intrinsic connectivity, all possible connections. (**b**) Family 1, input on pMTG. (**c**) Family 2, input on the bilateral pPHG. (**d**) Family 3 (winning): input on both pMTG and pPHG and their connections. The input was represented by High and Highest noise conditions (red triangle). Bayesian model averaging and Bayesian model selection criteria both surpassed a posterior probability of p>.99. L: left; R: right.
